# Atlas of ACE2 gene expression in mammals reveals novel insights in transmisson of SARS-Cov-2

**DOI:** 10.1101/2020.03.30.015644

**Authors:** Kun Sun, Liuqi Gu, Li Ma, Yunfeng Duan

**Affiliations:** Shenzhen Bay Laboratory, Shenzhen 518055, China; Beijing Huayuan Academy of Biotechnology, Beijing 100192, China

**Keywords:** COVID-19, 2019-nCov, novel coronavirus, potential host, susceptibility

## Abstract

**Background:** COVID-19 has become a worldwide pandemic. It is caused by a novel coronavirus named SARS-CoV-2 with elusive origin. SARS-CoV-2 infects mammalian cells by binding to ACE2, a transmembrane protein. Therefore, the conservation of ACE2 and its expression pattern across mammalian species, which are yet to be comprehensively investigated, may provide valuable insights into tracing potential hosts of SARS-CoV-2.

**Methods:** We analyzed gene conservation of ACE2 across mammals and collected more than 140 transcriptome datasets from human and common mammalian species, including presumed hosts of SARS-CoV-2 and other animals in close contact with humans. In order to enable comparisons across species and tissues, we used a unified pipeline to quantify and normalize ACE2 expression levels.

**Results:** We first found high conservation of ACE2 genes among common mammals at both DNA and peptide levels, suggesting that a broad range of mammalian species can potentially be the hosts of SARS-CoV-2. Next, we showed that high level of ACE2 expression in certain human tissues is consistent with clinical symptoms of COVID-19 patients. Furthermore, we observed that ACE2 expressed in a species-specific manner in the mammals examined. Notably, high expression in skin and eyes in cat and dog suggested that these animals may play roles in transmitting SARS-CoV-2 to humans.

**Conclusions:** Through building the first atlas of ACE2 expression in pets and livestock, we identified species and tissues susceptible to SARS-CoV-2 infection, yielding novel insights into the viral transmission.

## Introduction

In December 2019, outbreak of a novel coronavirus disease (COVID-19), a severe respiratory disease, and since spread rapidly throughout the country. A worldwide pandemic soon followed, with over 600,000 COVID-19 cases reported across 190 countries and territories by end of March 2020 and the number continues to rise sharply. After taking strong quarantine measures and national-wide lockdowns, the number of confirmed diagnoses is rapidly declining in China since February 2020. However, the endemic epicenter had since shifted to Italy and later to the United States. COVID-19 is becoming a global challenge to public health and continues to gather close attention.

The culprit of this pandemic is a new coronavirus named severe acute respiratory syndrome coronavirus 2 (SARS-CoV-2)^1^, which belongs to the same beta coronavirus family as SARS-CoV and MERS-CoV, the other two viruses that caused outbreaks in the past two decades^2^. Among myriad biological questions to be answered regarding SARS-CoV-2, one that is crucial and has intrigued much interest is the host origin and its mode of transmission to humans. SARS-CoV-2 and SARS-CoV are closely related, and both invade human cells via attaching their S proteins to a host transmembrane protein called angiotensin converting enzyme 2 (ACE2) as the entry point^3^. Using transcriptome data, some studies analyzed spatial expression patterns of ACE2 in various tissues and cell types of the human body, and reported that the receptor gene is indeed expressed in the lungs^4–6^. The connections between ACE2 expression and viral infection are further supported by cases from the United States, which confirmed for the first time the presence of SARS-CoV-2 in both the upper respiratory tract and stool sample of COVID-19 patients^7^.

Some studies have suggested that the original host of SARS-CoV-2 may be bats^2^. However, in the case of COVID-19, the outbreak occurred in winter when bats were under hibernation, making them unlikely to be the direct source of human infection. Hence, SARS-CoV-2 was likely transmitted to humans through some small carnivores like civet, the intermediate host of SARS-CoV. Recent studies have pointed to pangolins as the natural host of the virus, while minks being the possible intermediate host^8–10^. Virus tracing has thus been continuously carried out and attracted much attention. The mystery remains as which wild animals are definitive and intermediate hosts of the new coronavirus, and virus tracing has been continuously carried out and attracted much attention. Furthermore, is it possible that some of the animals living in close proximity to humans may also be susceptible to the virus and could potentially become additional hosts to SARS-CoV-2 hence further facilitating its transmission?

In attempt to address these questions, in this study we focused on the ACE2 gene, the host receptor of SARS-CoV-2 and other coronaviruses, and used quantitative data from various mammalian species to infer their susceptibility to SARS-CoV-2 infection. Since SARS-CoV-2 invade both bat and human cells via ACE2, we reasoned that if animals have ACE2 proteins similar to human, they could also become the targets of SARS-CoV-2, we thus evaluated the conservation of ACE2 gene across mammals. We further investigated ACE2 expression in various tissues among human and common mammals. In particular, we included species that live in close proximity with humans, i.e., pets and livestock. Our analyses identified potential species susceptible to SARS-CoV-2 and yielded novel insights into virus tracing and transmission, which may further contribute to the prevention and control of the COVID-19 pandemic.

## Methods

### Reference genomes and gene annotations

A total of 12 mammalian species were investigated in this study: *Homo sapiens* (human), *Mus musculus* (mouse), *Rhinolophus sinicus* (Chinese rufous horseshoe bat), *Manis javanica* (pangolin), *Felis catus* (cat), *Canis lupus familiaris* (dog), *Mustela putorius furo* (ferret), *Mesocricetus auratus* (hamster), *Bos taurus* (cow), *Sus scrofa* (pig), *Oryctolagus cuniculus* (rabbit), and *Capra hircus* (goat). Among these mammals, bats and pangolins are presumed candidates of definitive and intermediate hosts of SARS-CoV-2, respectively^2,11^; cats, dogs, ferrets and hamsters are common pets; cows, pigs, rabbits and goats are common livestock. Latest versions of reference genomes and RefSeq gene annotations^12^ for these species were downloaded from NCBI (National Center for Biotechnology Information)^13^. Detailed information was described in Table S1 in Supplementary data.

### Transcriptome data collection

Transcriptome data for various mammalian tissues generated from 142 RNA-seq experiments (whole transcriptome shotgun sequencing) was collected from publicly available sources. Briefly, human data was from the GTEx (Genotype-Tissue Expression) project^14^, ENCODE (ENCyclopedia Of DNA Elements) project^15^, and Frausto et al.^16^; bat data was from Dong et al.^17^; pangolin data was from Ma et al.^18^ and Mohamed Yusoff et al.^19^; cat data was from 99 Lives Cat Genome Sequencing Initiative project; dog data was from Hoeppner et al.^20^, Lindblad-Toh et al.^21^ and Sudharsan et al.^22^; ferret and rabbit data was from Chen et al.^23^; hamster and goat data was from Fushan et al.^24^; cow data was from Merkin et al.^25^; and pig data was from Summers et al.^26^. Detailed information on accession numbers and tissues for each species was listed in Table S2 in Supplementary data.

### Transcriptome data analysis

All the transcriptome data was analyzed using a unified pipeline. Briefly, raw RNA-seq reads were first preprocessed to trim sequencing adapters and low-quality cycles using Ktrim software^27^ with default parameters; the preprocessed reads were then aligned to corresponding reference genomes using STAR software^28^ with default parameters. Key statistics during preprocessing and alignment was present in Table S2 in Supplementary data. The vast majority (141/142) of the samples had more than 10 million uniquely mapped reads (median: 27.8 million), indicating sufficient sequencing depths for reliable gene expression quantifications^29^, which was performed using featureCounts^30^ software with default parameters against RefSeq gene annotations^12^. Considering that the reference genomes and gene annotations for most of the species included in this study are far from complete, in order to avoid potential biases of the conventional FPKM (Fragments per Kilobase Million) values, we used ACTB (Actin Beta) gene from each RNA-seq experiment to normalize ACE2 expression for appropriate comparisons across species and tissue types. ACTB is a housekeeping gene that is abundantly and stably expressed in most cell types, and is commonly used as an internal control for gene expression normalizations^31^. In addition, ACTB gene is also conserved in all the species investigated in this study. The following formula was used to calculate normalized ACE2 expression:

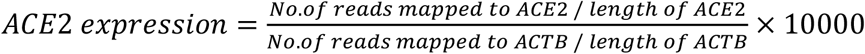

We used 10000 as a scale factor here because ACE2 was typically expressed at much lower levels than ACTB. Meanwhile, since data from the GTEx project was provided as preprocessed values, these GTEx values were used directly in the above formula in lieu of number of mapped reads and gene length.

## Results

### Conservation of ACE2 gene in mammals

We first examined the conservation of ACE2 across mammalian species. A snapshot obtained from the UCSC genome browser^32^ was shown in Figure 1A, which demonstrated high conservation of ACE2 gene’s exons in DNA level. Next we examined the conservation of ACE2 at the peptide level by searching the UniProt database^33^ for ACE2 protein sequences in mammals. We limited the analysis to species listed in Methods (plus rats) and discarded duplicate records as well as records with abnormally short sequences (< 300 amino acids), leaving 24 ACE2 protein sequences for subsequent analyses. Pairwise alignment results showed that 23 non-human ACE2 proteins share high level of similarity to the human ACE2 protein, with the identity scores ranging from 78.6% to 85.2% and a median score of 81.9%. Notably, for the two virus-binding hotspots (i.e., the 31th and 353th amino acid)^34^, all the mammals except mice and rats share the same amino acids to the human ACE2 protein (Table S3 in Supplementary data). Surprisingly, in contrast to conventional phylogenetic tree generated using genomic data, the phylogenetic tree based on the ACE2 proteins showed that, cats and dogs were the species closest to human instead of mice among the mammals included in this analysis (Figure 1B). Taken together, the conservation analysis showed that ACE2 gene was highly conserved among common mammals at both DNA and peptide levels, suggesting that SARS-CoV-2 can potentially bind to ACE2 proteins in these mammals with high affinity^2^.

**Figure 1.**
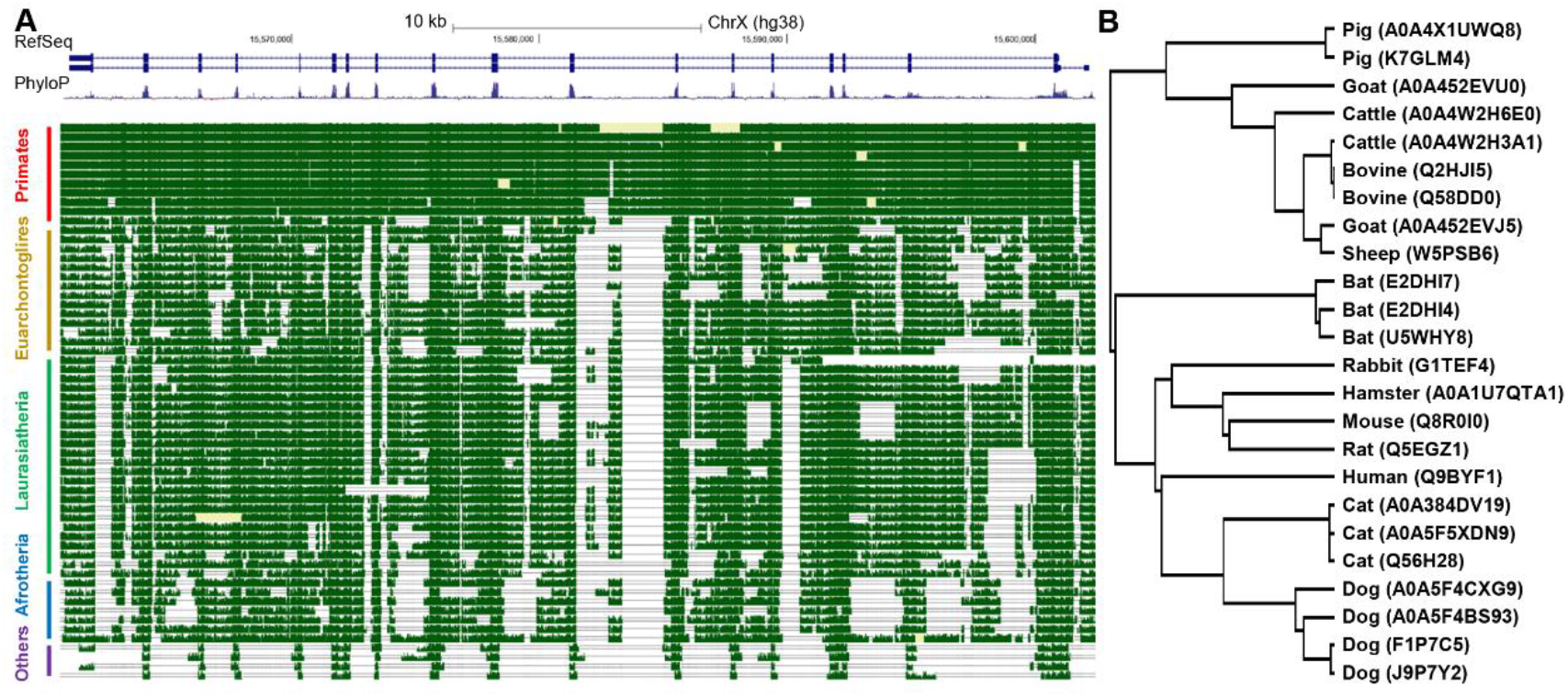
Conservation analysis of ACE2 gene. (A) Snapshot from the UCSC genome browser showing the DNA level conservation of ACE2 gene. Note that horse was removed due to lack of pair-wise alignment data. (B) Phylogenetic tree based on mammal ACE2 protein sequences from UniProt database. The top panel in (A) is RefSeq annotation of ACE2 gene in human (which contains two isoforms), the middle panel is conservation scores among 100 vertebrates, higher values denote higher levels of conservation), and the best-in-genome pairwise alignments among 42 mammals are plotted in the bottom panel. For (B), the protein accession number in UniProt were shown in parentheses and some species has multiple records.

### Expression profile of ACE2 gene in human tissues

We then profiled the expression patterns of ACE2 gene in human tissues. Data from the GTEx project showed that ACE2 was expressed in various tissues, including testis, intestines, heart, kidney, and pancreas (Figure 2A). It was worth noting that our normalized ACE2 expression pattern was similar to the original version obtained from the GTEx portal (Figure S1 in Supplementary data), except that our analysis highlighted the heart as the tissue with 3rd highest ACE2 expression.

**Figure 2.**
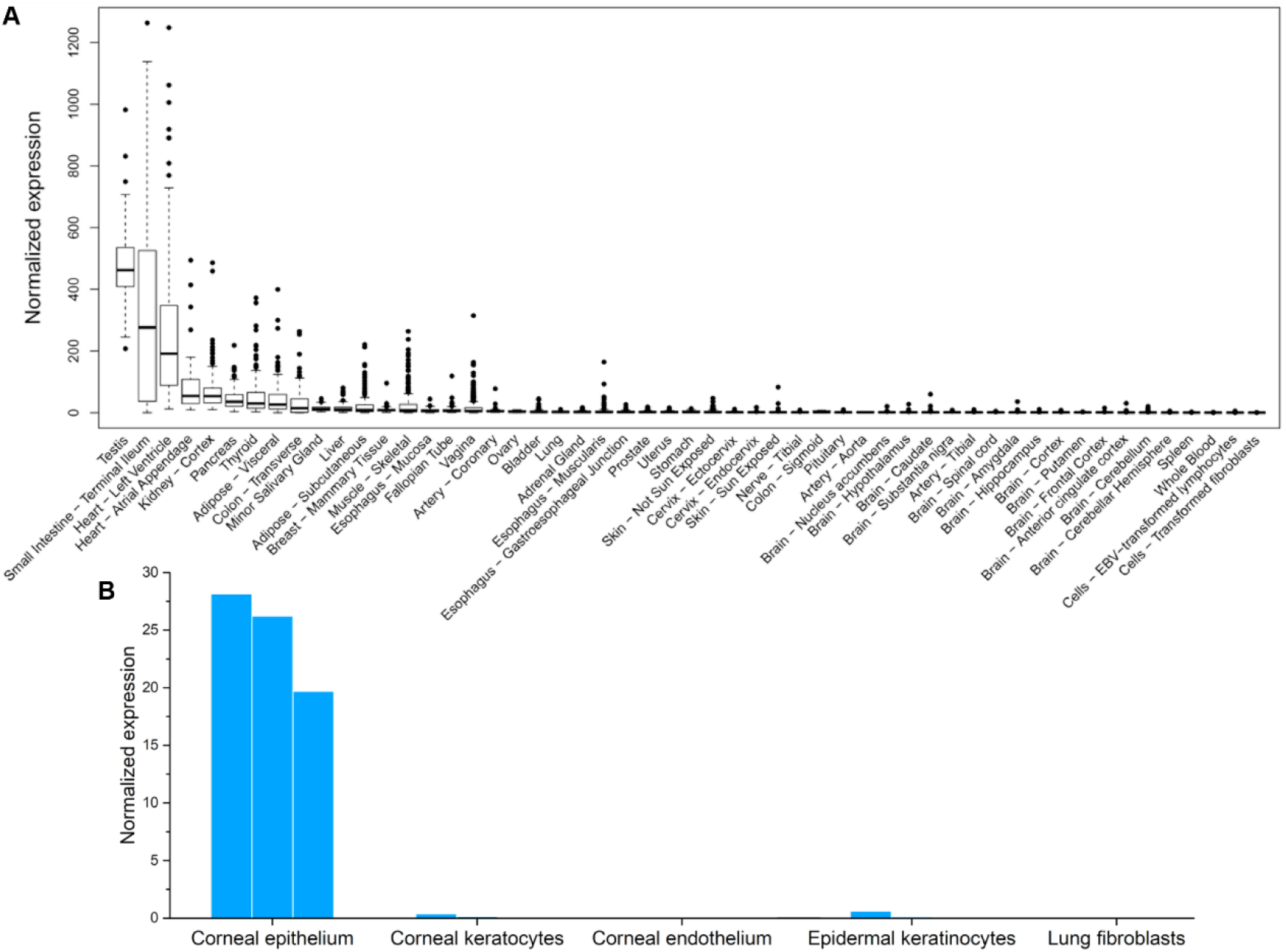
Expression of ACE2 gene in human tissues. Data was collected from (A) GTEx project, (B) corneal tissues, epidermal keratinocytes, lung fibroblasts. Replicate experiments for tissues in panel B were available and were shown as multiple bars.

We also collected transcriptome data from tissues that were not in the GTEx datasets, including cornea, epidermal keratinocytes, and lung fibroblasts. These tissues are frequently exposed to the air. ACE2 gene was expressed in corneal epithelial cells (Figure 2B), suggesting that the eye could be vulnerable to SARS-CoV-2 infections. In contrast, ACE2 was not expressed in epidermal keratinocytes nor lung fibroblasts, which was consistent with the above analysis based on GTEx data. This result was also consistent with a previous report based on single-cell RNA-seq data that ACE2 was only expressed in the alveolar type 2 (AT2) cells in lungs^35^.

### Expression pattern of ACE2 gene in mice

Mouse is the most widely used model species in biomedical studies, including those related to SARS-CoV-2. We extracted the expression data of murine Ace2 gene from Tabula Muris project^36^, which investigated various murine tissues using single-cell RNA-seq experiments. Murine Ace2 gene was expressed in kidney, heart, intestine, and pancreas (Figures 3 and S1 in Supplementary data), a pattern similar to human (Figure 2A). However, murine Ace2 gene was not expressed in any cell types in lungs, while expressed in tongue and skin (Figure 3B). The data also suggested that ACE2 gene expression pattern could be species-specific among mammals. In particular, lung-related symptoms may not be expected when infecting SARS-CoV-2 to normal mice.

**Figure 3.**
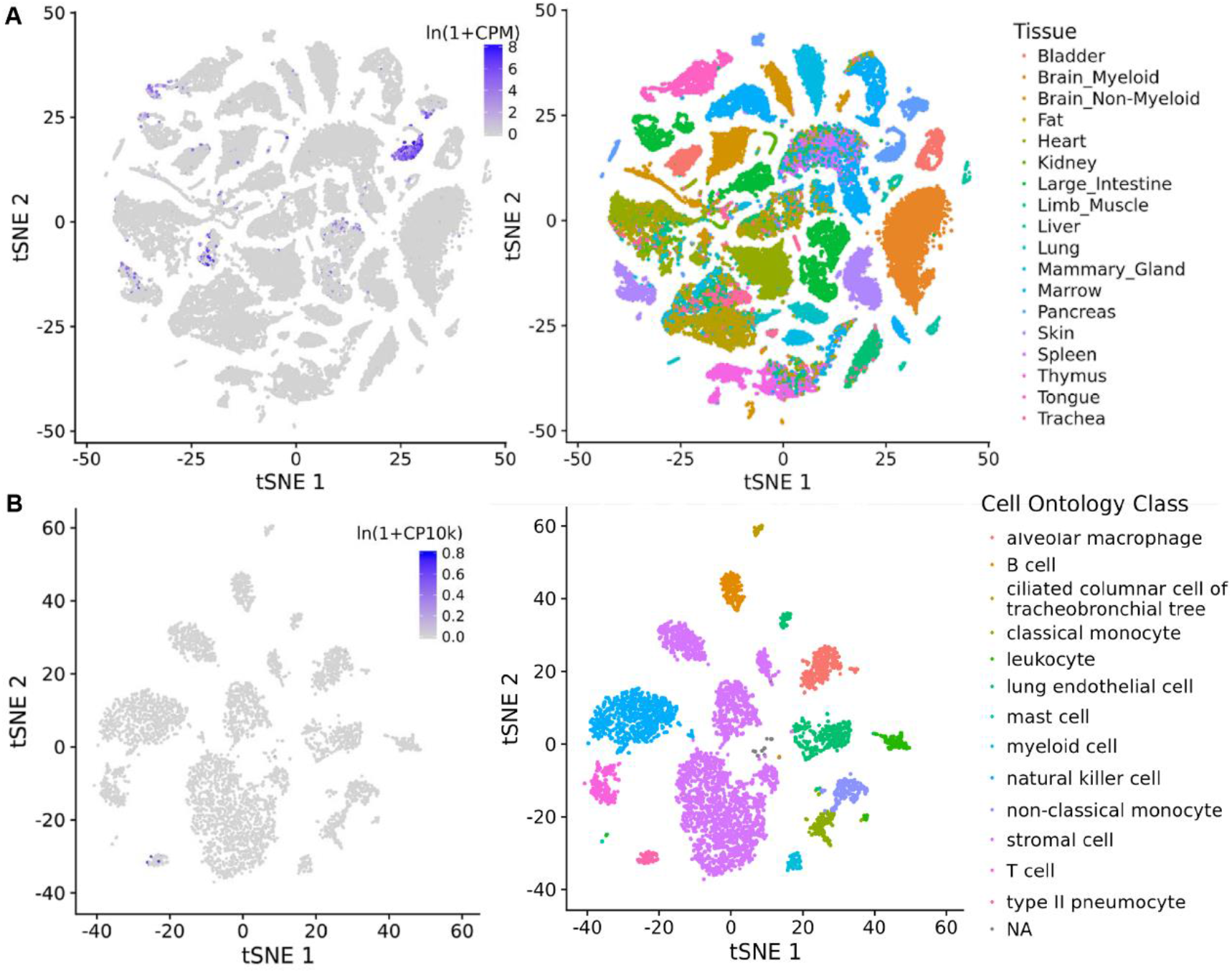
Expression of ACE2 in murine tissues. Data was obtained from the Tabula Muris project. (A) Expression in all cells isolated using FACS protocol (left) and corresponding tissue origin of these cells (right). Note that the cells were clustered using t-SNE (t-distributed Stochastic Neighbor Embedding) algorithm based on their transcriptome. (B) Expression in lung tissue.

### Expression patterns of ACE2 gene in other mammals

Bats and pangolins are hypothesized to be the definitive and intermediate hosts of SARS-Cov-2^2^. We profiled the expression of ACE2 in various tissues in Chinese rufous horseshoe bats and Malayan pangolins. ACE2 gene was highly expressed in most tissues examined in both bats and in pangolin, including those frequently exposed to the air, e.g., lungs in bats and tongue in pangolins (Figure 4).

**Figure 4.**
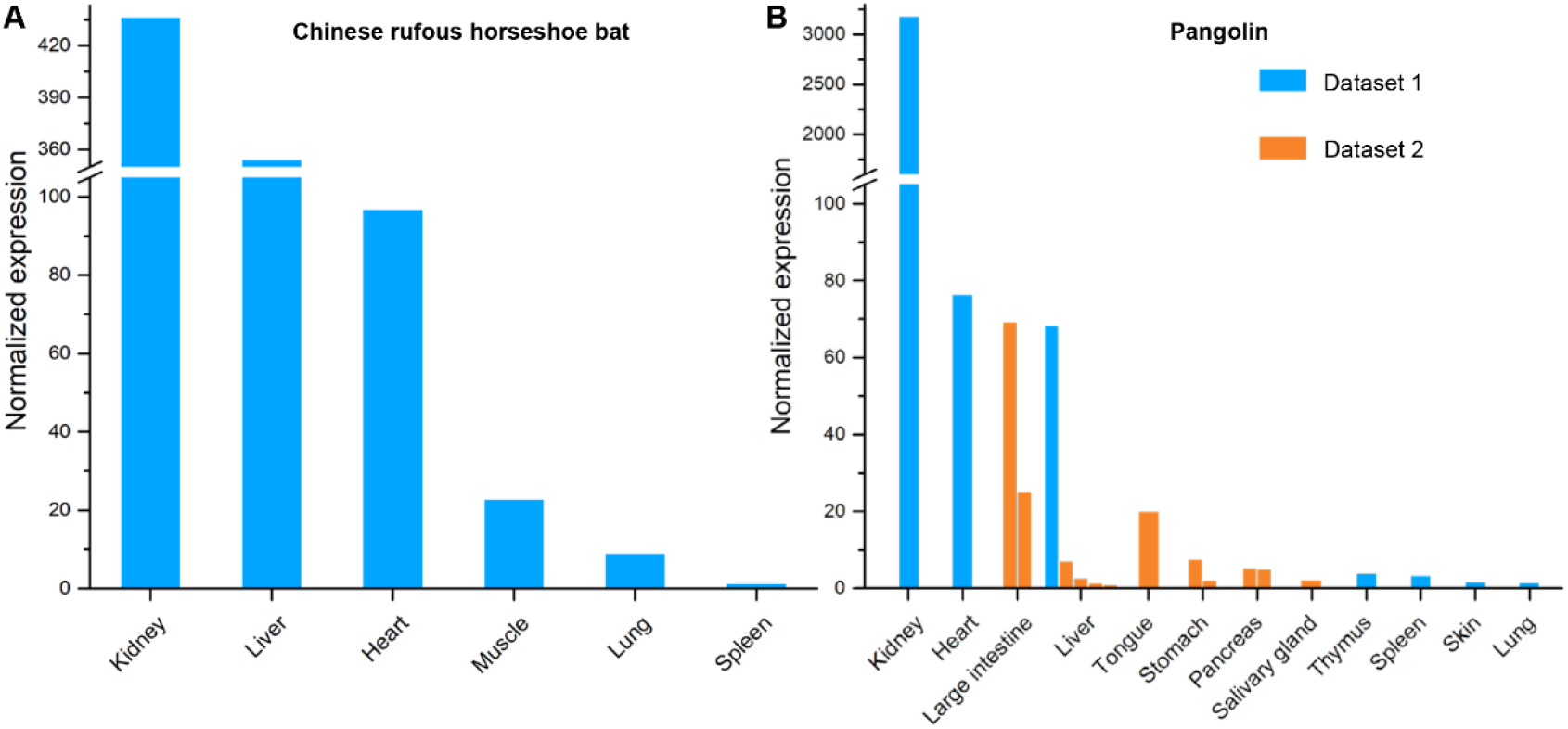
Expression of ACE2 gene in (A) bat and (B) pangolin tissues. Two datasets for pangolin tissues were collected and labelled with different colors. Replicate experiments for some tissues were available and were shown as multiple bars.

Pets are the most intimate animals to humans and may thus very likely to be contracted by human COVID-19 patients or to transmit the virus to humans if they are infected. We examined ACE2 expression patterns in cats and dogs, the most popular pets worldwide, as well as ferrets and hamsters, which are also very common in China. ACE2 gene was highly expressed in various tissues in these animals, such as kidney, heart and liver (Figures 5A-5C and S2A in Supplementary data). For cats, ACE2 was also highly expressed in skin, ear tip, lungs, and retina; for dogs, ACE2 was expressed in skin and retina. This data suggested that cats and dogs may be highly susceptible to SARS-CoV-2 infection. In addition, we also observed ACE2 expression in the lungs of cats and ferrets, which suggested that these animals may be more suitable for SARS-CoV-2 studies than rodent models^2,37^.

**Figure 5.**
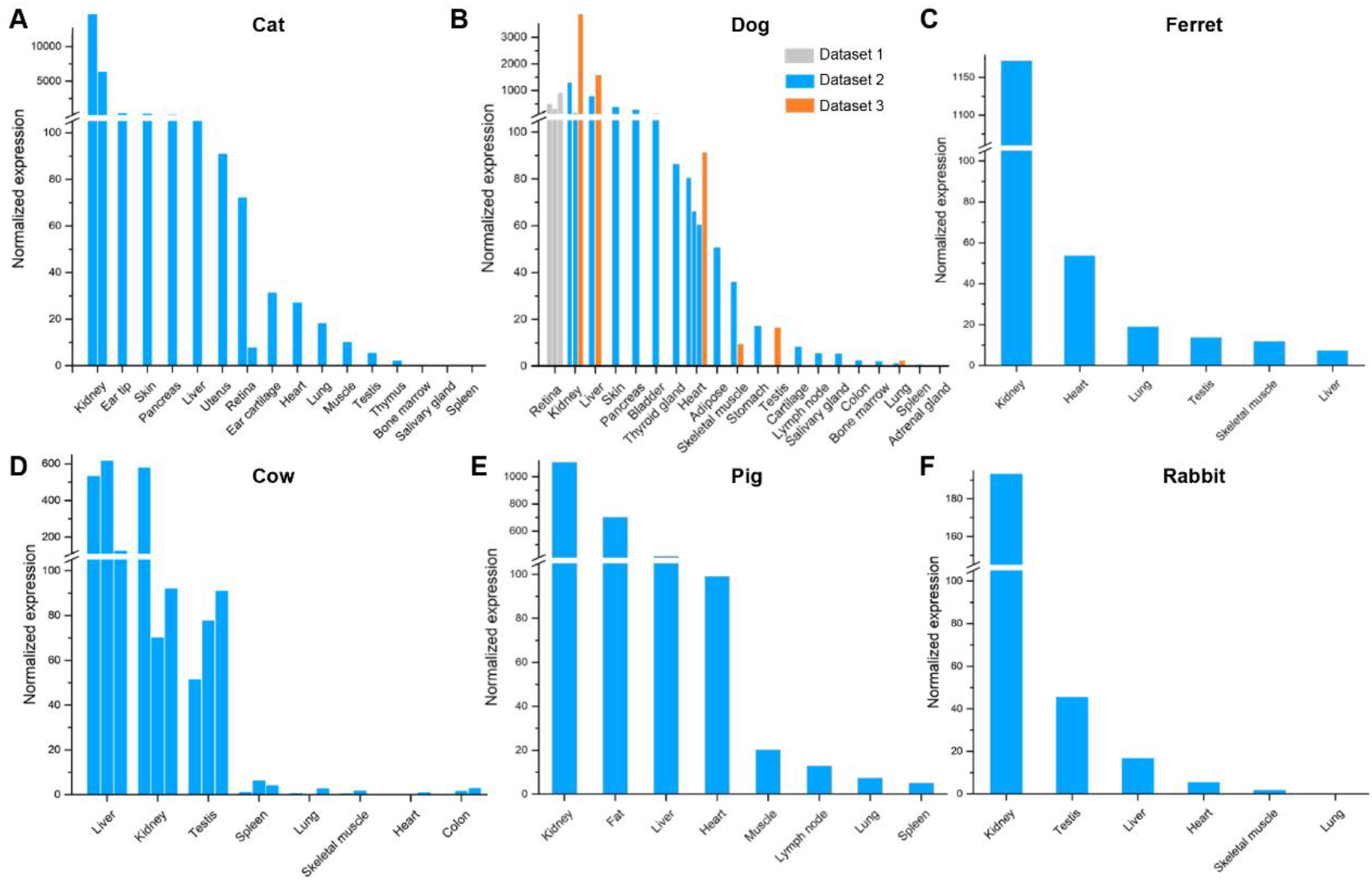
Expression of ACE2 in (A) cat, (B) dog, (C) ferret, (D) pig, (E) cow and (F) rabbit tissues. Three datasets for dog tissues were collected and labelled with different colors. Replicate experiments for some tissues were available and were shown as multiple bars.

Besides pets, livestock are another category of mammals that are frequently in direct contact with humans. We profiled the ACE2 expression in cows, pigs, rabbits (which can also serve as pets), and goats. In these animals, ACE2 was highly expressed in multiple tissues including kidney, liver, and heart, but not in muscles (Figures 5D-5F and S2B in Supplementary data). ACE2 was also highly expressed in the fat of pigs. The data suggested that storage of fresh or undercooked meat, especially viscera tissues, might be risky.

## Discussion

We showed that ACE2 expression profile in human tissues was in agreement with clinical observations of SARS-CoV-2 infected individuals. For example, ACE2 was by far most strongly expressed in testis. SARS is known to cause orchitis and destruction to male germ cells^38^. Similar effect from COVID-19 was speculated, but a most recent study had just provided the first direct evidence that SARS-CoV-2 infection impacts male sex hormones^39^. Similarly, small intestine is the tissue with the second highest level of ACE2 expression, while SARS-CoV-2 was found in stool samples of COVID-19 patients both in China and the United States^7,42^, even after the viral RNA decreased to undetectable level in respiratory tract^43^. Additionally, we found in our normalized dataset that heart tissues also had the third highest level of ACE2 expression. This observation is also in line with reports showing that COVID-19 patients complicated with cardiac diseases are subject to the highest mortality risk^40,41^. Together, the consistence between our results and clinical observations demonstrated that ACE2 expression is a reasonable indicator for susceptibility to SARS-CoV-2 infection.

Our analyses showed that mammalian ACE2 genes are highly conserved across lineages and exhibit broad expression patterns. However, tissue-specific expression profiles varied from species to species, suggesting certain mammals may be the hosts of SARS-CoV-2, i.e., carriers without significant symptoms. Compared to humans, other mammals examined in this study either don’t express ACE2 in their testes, or at a low level. On the contrary, among non-human species kidney was almost always the tissue with highest ACE2 expression. Meanwhile, it is worth noting that mouse is a common model species for many human medical studies, including COVID-19. However, we showed that in ACE2 proteins, mice and rats are the only mammals that have different amino acids in the two virus-binding hotspots from humans, which is consistent with previous *in vitro* virus infection experiments^2^. Moreover, ACE2 was barely expressed in any of the cell types in murine lungs. Thus, both observations argue against using mice as the optimal animal model for studying coronavirus related diseases.

Most notably, our analyses revealed that ACE2 expression levels are particularly high in cats and dogs. Especially in cats, expression levels in top four ACE2 expression hotspot tissues are all magnitudes higher than any other mammals examined. Intriguingly, in the ACE2 phylogenic tree, both cats and dogs are also cluster closer to humans than other mammals (Figure 1B), suggesting potentially higher affinity to coronaviruses in these pets. SARS-CoV is known to infect cats^44^, while SARS-CoV-2 positive cats and dogs had also been reported^45,46^, therefore our analyses suggest high possibility that cats and dogs can host SARS-CoV-2. Furthermore, in both cats and dogs, skin had high abundant ACE2 expression, so did ear tip in cats and retina in dogs. High ACE2 expression levels in these exterior body parts makes them particularly likely to host SARS-CoV-2 and pass on to humans via skin to skin contact. Stray animals can even be more serious transmitters of coronaviruses. It is estimated that there are approximately 500 million stray dogs and similar number of stray cats worldwide^47^. Cats and dogs are sometimes slaughtered for meat, including a large proportion of stray ones^48^. Therefore, it is likely that cats and dogs may have contributed to the COVID-19 pandemic. For the least, people should also be vigilant with handling of pets in the effort of containing the spread of SARS-CoV-2.

In conclusion, in this study, we had reported the high conservation and built the first expression atlas of ACE2 gene in common mammals. Our analyses revealed species and tissues susceptible to SARS-CoV-2 infection, thus yielding novel insights in tracing the origin and transmission of the virus.

## Acknowledgements

This work has been supported by Shenzhen Bay Laboratory and Beijing Huayuan Academy of Biotechnology.

