## Supplementary materials for "Atlas of ACE2 gene expression in mammals reveals novel insights in transmisson of SARS-Cov-2"

### Table of Contents

**Figure S1.** Expression of ACE2 in human tissues. ---------------------------------------- 2

**Figure S2.** Expression of ACE2 in murine tissues. ---------------------------------------- 3

**Figure S3.** Normalized expression of ACE2 in hamsters and goats. --------------------- 4

**Table S1.** Reference genomes and annotations used in this study. ---------------------- 5

**Table S2.** Accession and statistics of the transcriptome data compiled in this study. -- 6

**Table S3.** Alignment result of mammal ACE2 protein sequences in UniProt. ---------- 9

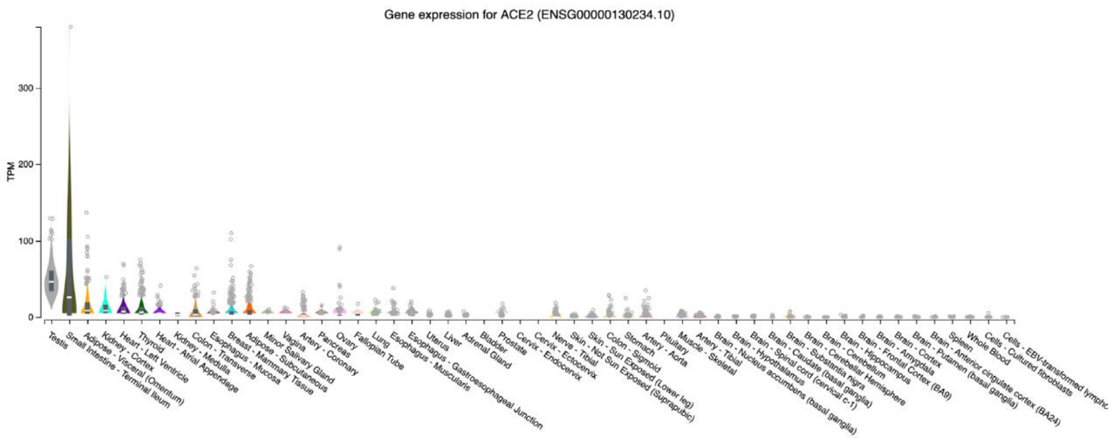


**Figure S1.** Expression of ACE2 in human tissues. The figure was obtained from GTEx portal.

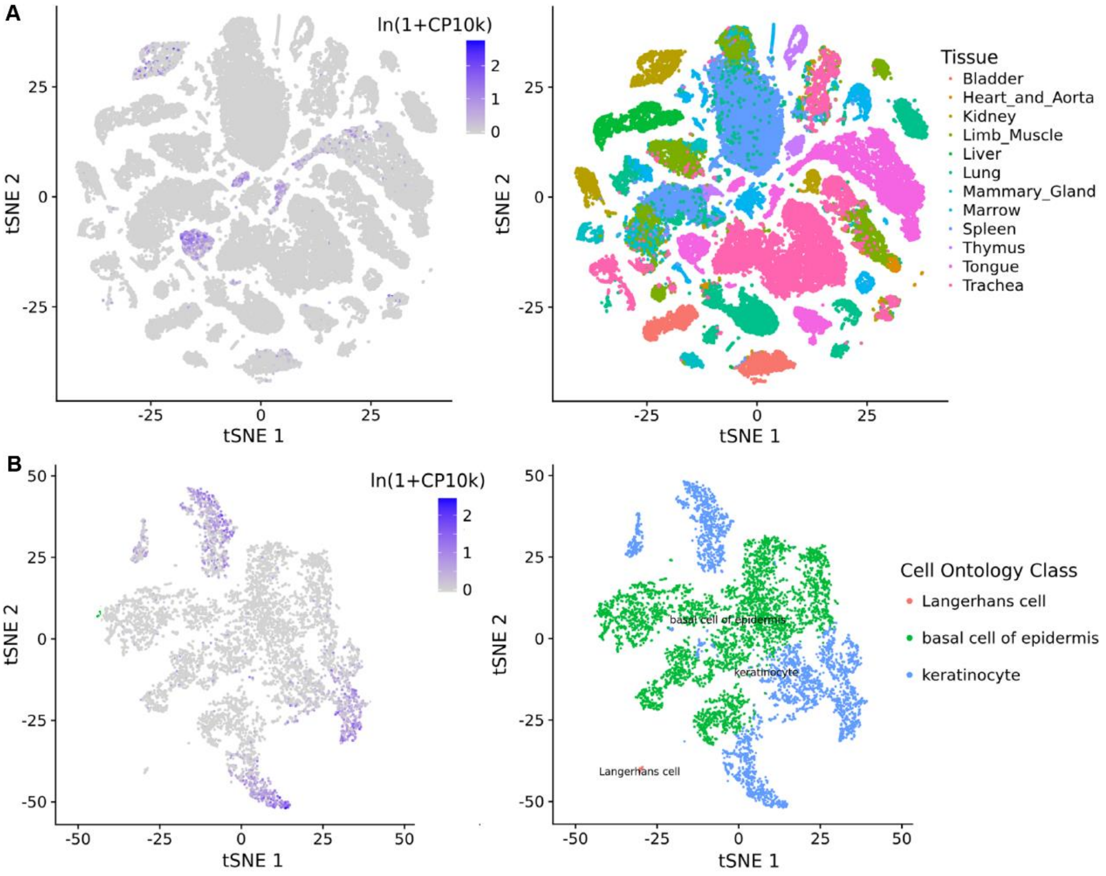


**Figure S2.** Expression of ACE2 in murine tissues. The data was obtained from Tabula Muris project. (A) All tissues. (B) Tongue.

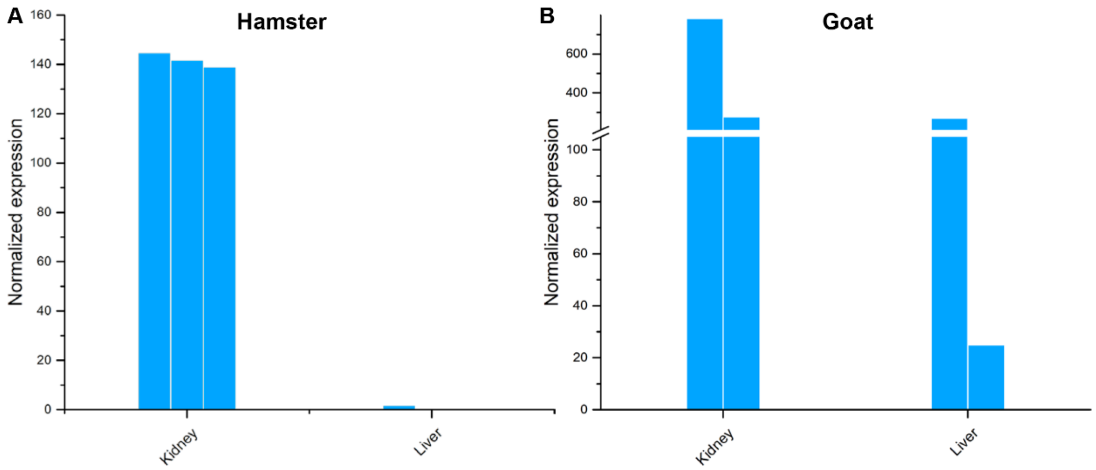


**Figure S3.** Expression of ACE2 in (A) hamster and (B) goat tissues.

**Table S1. Reference genomes and gene annotation used in this study.**

| Species | Common name | Assembly name | Assembly accession | Annotation release ID |
|---|---|---|---|---|
| Homo sapiens | Human | GRCh38.p13 | GCF_000001405.39 | 109 |
| Rhinolophus sinicus | Bat | ASM188883v1 | GCF_001888835.1 | 100 |
| Manis javanica | Manis | ManJav1.0 | GCF_001685135.1 | 100 |
| Felis catus | Cat | Felis_catus_9.0 | GCF_000181335.3 | 104 |
| Canis lupus familiaris | Dog | CanFam3.1 | GCF_000002285.3 | 105 |
| Mustela putorius furo | Ferret | MusPutFur1.0 | GCF_000215625.1 | 101 |
| Mesocricetus auratus | Hamster | MesAur1.0 | GCF_000349665.1 | 102 |
| Bos taurus | Cow | ARS-UCD1.2 | GCF_002263795.1 | 106 |
| Sus scrofa | Pig | Sscrofa11.1 | GCF_000003025.6 | 106 |
| Oryctolagus cuniculus | Rabbit | OryCun2.0 | GCF_000003625.3 | 102 |
| Capra hircus | Goat | ARS1 | GCF_001704415.1 | 102 |

### Table S2. Accession and statistics of the transcriptome data compiled in this study.

| Species | Data source | Sample ID | Tissue information | Total reads | Ktrim preprocessed | STAR aligned | Mappa-bility(%) |
|---|---|---|---|---|---|---|---|
| Homo sapiens | ENCODE | NHEK.rep1 | Epidermal | 68,932,471 | 42,944,842 | 36,715,777 | 81.9 |
| | | NHEK.rep2 | Epidermal | 68,995,645 | 60,849,039 | 58,067,559 | 89.2 |
| | | NHLF.rep1 | Lung fibroblasts | 67,321,664 | 59,671,441 | 56,028,139 | 88.5 |
| | | NHLF.rep2 | Lung fibroblasts | 67,164,915 | 55,872,307 | 50,225,300 | 89.0 |
| | GSE121922 | SRR8129827 | Corneal epithelium | 21,384,188 | 21,063,173 | 17,912,697 | 85.0 |
| | | SRR8129828 | Corneal epithelium | 20,913,971 | 20,625,789 | 17,502,545 | 84.9 |
| | | SRR8129829 | Corneal epithelium | 19,466,412 | 19,187,323 | 16,184,315 | 84.4 |
| | | SRR8129830 | Corneal keratocytes | 21,730,275 | 21,495,198 | 18,156,122 | 84.5 |
| | | SRR8129831 | Corneal keratocytes | 18,799,079 | 18,527,115 | 15,575,797 | 84.1 |
| | | SRR8129832 | Corneal keratocytes | 18,996,222 | 18,718,811 | 15,701,517 | 83.9 |
| | | SRR8129833 | Corneal endothelium | 13,389,782 | 13,138,066 | 10,564,327 | 80.4 |
| | | SRR8129834 | Corneal endothelium | 12,285,592 | 12,019,769 | 9,653,549 | 80.3 |
| | | SRR8129835 | Corneal endothelium | 13,928,169 | 13,644,250 | 10,959,195 | 80.3 |
| Rhinolophus sinicus | SRP063381 | SRR2273931 | Kidney | 30,337,622 | 29,795,046 | 26,698,695 | 89.6 |
| | | SRR2273875 | Liver | 30,559,494 | 30,057,890 | 27,893,166 | 92.8 |
| | | SRR2273739 | Heart | 27,788,814 | 27,274,122 | 22,964,399 | 84.2 |
| | | SRR2273816 | Muscle | 32,875,990 | 32,299,770 | 25,715,822 | 79.6 |
| | | SRR2273738 | Brain | 33,101,848 | 32,507,454 | 30,637,988 | 94.3 |
| | | SRR2273740 | Lung | 34,488,781 | 33,837,363 | 32,558,189 | 96.2 |
| | | SRR2273762 | Spleen | 22,008,969 | 20,889,127 | 11,166,880 | 53.5 |
| Manis javanica | SRP156258 | SRR7641079 | Pancreas | 24,856,498 | 24,856,472 | 20,906,101 | 84.1 |
| | | SRR7641080 | Liver | 21,967,580 | 21,967,553 | 18,689,870 | 85.1 |
| | | SRR7641081 | Liver | 23,980,609 | 23,980,571 | 19,980,166 | 83.3 |
| | | SRR7641082 | Pancreas | 24,376,771 | 24,376,734 | 20,839,958 | 85.5 |
| | | SRR7641083 | Tongue | 28,411,058 | 28,411,029 | 24,114,077 | 84.9 |
| | | SRR7641084 | Salivary glands | 22,444,882 | 22,444,842 | 19,046,428 | 84.9 |
| | | SRR7641085 | Stomach | 22,856,344 | 22,856,313 | 19,682,763 | 86.1 |
| | | SRR7641086 | Stomach | 25,508,428 | 25,508,397 | 21,964,813 | 86.1 |
| | | SRR7641087 | Liver | 25,805,479 | 25,805,442 | 21,541,960 | 83.5 |
| | | SRR7641088 | Liver | 28,768,081 | 28,768,050 | 24,409,509 | 84.9 |
| | | SRR7641089 | Large Intestine | 29,573,813 | 29,573,794 | 25,137,147 | 85.0 |
| | | SRR7641090 | Large Intestine | 20,449,264 | 20,449,253 | 18,203,703 | 89.0 |
| | SRP064341 | SRR3923846 | Skin | 24,805,754 | 24,493,823 | 21,702,376 | 88.6 |
| | | SRR2561214 | Lungs | 45,883,537 | 45,883,536 | 43,005,271 | 93.7 |
| | | SRR2561215 | Spleen | 52,632,086 | 52,632,086 | 49,422,337 | 93.9 |
| | | SRR2561213 | Liver | 45,306,423 | 45,306,423 | 42,722,562 | 94.3 |
| | | SRR2561216 | Thymus | 41,532,741 | 41,532,741 | 39,554,501 | 95.2 |
| | | SRR2561212 | Kidney | 42,419,462 | 42,419,462 | 40,640,322 | 95.8 |
| | | SRR2561211 | Heart | 47,079,243 | 47,079,243 | 45,497,321 | 96.6 |
| Felis catus | SRP071078 | SRR3200449 | Lung | 61,604,161 | 59,373,190 | 52,323,679 | 88.1 |
| | | SRR3200451 | Muscle | 53,498,415 | 51,628,230 | 44,553,838 | 86.3 |
| | | SRR3200453 | Liver | 66,486,821 | 64,279,254 | 54,290,629 | 84.5 |
| | | SRR3200454 | Ear Tip | 61,849,569 | 59,863,742 | 49,458,043 | 82.6 |
| | | SRR3200455 | Ear cartilage | 74,477,894 | 72,375,687 | 67,892,280 | 93.8 |

| | | | | | | | |
|---|---|---|---|---|---|---|---|
| | | SRR3200457 | Retina | 54,310,271 | 52,850,253 | 46,389,760 | 87.8 |
| | | SRR3200458 | Uterus | 51,634,724 | 48,975,318 | 40,155,738 | 82.0 |
| | | SRR3200459 | Bone marrow | 36,096,745 | 33,351,045 | 24,701,766 | 74.1 |
| | | SRR3200460 | Kidney | 60,295,175 | 58,082,781 | 43,644,190 | 75.1 |
| | | SRR3200462 | Testes | 80,274,030 | 77,645,779 | 70,923,239 | 91.3 |
| | | SRR3200465 | Retina | 58,621,360 | 57,043,848 | 51,114,268 | 89.6 |
| | | SRR3200469 | Pancreas | 83,078,528 | 80,123,579 | 74,509,518 | 93.0 |
| | | SRR3200470 | Skin | 66,777,859 | 64,602,457 | 55,718,935 | 86.3 |
| | | SRR3200471 | Heart | 107,724,558 | 104,532,937 | 67,788,907 | 64.9 |
| | | SRR3200473 | Kidney | 47,219,252 | 45,690,466 | 34,964,638 | 76.5 |
| | | SRR3218714 | Spleen | 60,725,845 | 58,500,972 | 49,693,589 | 84.9 |
| | | SRR3218715 | Thymus | 52,195,042 | 49,203,820 | 41,479,120 | 84.3 |
| | | SRR3218716 | Spleen | 42,845,814 | 41,577,135 | 34,374,919 | 82.7 |
| | | SRR3218717 | Salivary gland | 56,827,581 | 54,141,092 | 45,207,119 | 83.5 |
| Canis lupus familiaris | GSE97638 | SRR5443379 | Neural retina | 18,158,136 | 18,138,066 | 17,449,363 | 96.2 |
| | | SRR5443380 | Neural retina | 25,489,581 | 25,465,359 | 24,490,231 | 96.2 |
| | | SRR5443381 | Neural retina | 12,003,881 | 11,992,690 | 11,535,499 | 96.2 |
| | SRP114662 | SRR8996983 | Adipose | 45,152,409 | 43,618,974 | 40,633,451 | 93.2 |
| | | SRR8996995 | Adrenal gland | 55,999,077 | 54,435,675 | 42,867,568 | 78.8 |
| | | SRR8996996 | Bladder | 25,629,091 | 24,710,475 | 23,381,661 | 94.6 |
| | | SRR8996993 | Bone marrow | 72,272,259 | 69,830,421 | 59,939,651 | 85.8 |
| | | SRR8996994 | Cartilage | 53,915,899 | 52,097,984 | 49,142,410 | 94.3 |
| | | SRR8997000 | Colon | 44,721,489 | 43,036,543 | 40,558,438 | 94.2 |
| | | SRR8997001 | Kidney (cortex) | 57,787,581 | 56,189,631 | 51,652,382 | 91.9 |
| | | SRR8997002 | Kidney (medulla) | 44,087,155 | 42,677,016 | 39,643,305 | 92.9 |
| | | SRR8996964 | Heart (left atrium) | 46,569,235 | 45,213,751 | 42,470,278 | 93.9 |
| | | SRR8996963 | Heart (left ventricle) | 45,991,318 | 44,449,475 | 42,594,327 | 95.8 |
| | | SRR8996969 | Heart (right ventricle) | 46,648,476 | 45,117,648 | 42,915,864 | 95.1 |
| | | SRR8996966 | Liver | 51,692,060 | 49,953,096 | 46,598,407 | 93.3 |
| | | SRR8996965 | Lung | 46,168,334 | 44,590,607 | 40,816,349 | 91.5 |
| | | SRR8996968 | Lymph node | 62,480,995 | 60,629,005 | 51,758,660 | 85.4 |
| | | SRR8996967 | Pancreas | 55,283,372 | 53,379,088 | 46,388,546 | 86.9 |
| | | SRR8996972 | Salivary gland | 43,256,190 | 41,697,927 | 36,926,974 | 88.6 |
| | | SRR8996971 | Skeletal muscle | 45,631,797 | 44,233,179 | 41,948,357 | 94.8 |
| | | SRR8997043 | Skin | 43,200,562 | 41,534,641 | 38,317,594 | 92.3 |
| | | SRR8997044 | Spleen | 26,950,137 | 26,050,311 | 21,795,307 | 83.7 |
| | | SRR8997045 | Stomach | 55,518,261 | 53,647,070 | 50,276,470 | 93.7 |
| | | SRR8997046 | Thyroid gland | 48,423,432 | 46,874,668 | 43,968,733 | 93.8 |
| | GSE97638 | SRR6206901 | Heart | 31,934,896 | 28,297,899 | 25,005,163 | 88.4 |
| | | SRR6206906 | Kidney | 35,927,711 | 31,403,005 | 27,697,606 | 88.2 |
| | | SRR6206911 | Liver | 33,858,041 | 30,213,606 | 23,632,522 | 78.2 |
| | | SRR6206916 | Lung | 29,502,542 | 25,638,471 | 21,877,766 | 85.3 |
| | | SRR6206921 | Skeletal Muscle | 32,712,735 | 28,511,422 | 24,698,601 | 86.6 |
| | | SRR6206926 | Testis | 35,435,315 | 30,667,393 | 26,339,541 | 85.9 |
| Mustela putorius furo | GSE106077 | SRR6206902 | Heart | 29,394,130 | 27,861,205 | 26,291,047 | 94.4 |
| | | SRR6206907 | Kidney | 27,472,302 | 25,711,807 | 24,206,747 | 94.2 |
| | | SRR6206912 | Liver | 27,454,811 | 25,996,290 | 24,282,703 | 93.4 |
| | | SRR6206917 | Lung | 26,849,508 | 25,065,001 | 23,473,250 | 93.7 |
| | | SRR6206922 | Skeletal muscle | 30,505,719 | 28,822,176 | 27,180,993 | 94.3 |

| | | | | | | | |
|---|---|---|---|---|---|---|---|
| | | SRR6206927 | Testis | 27,024,467 | 25,551,148 | 23,846,337 | 93.3 |
| Mesocricetus auratus | GSE43013 | SRR636857 | Liver | 22,166,056 | 21,444,610 | 19,015,456 | 88.7 |
| | | SRR636858 | Liver | 15,859,602 | 15,549,003 | 13,966,154 | 89.8 |
| | | SRR636859 | Liver | 19,789,823 | 19,142,756 | 16,996,355 | 88.8 |
| | | SRR636907 | Kidney | 21,084,573 | 20,327,600 | 18,417,620 | 90.6 |
| | | SRR636908 | Kidney | 22,131,879 | 21,703,453 | 19,566,391 | 90.2 |
| | | SRR636909 | Kidney | 16,437,102 | 16,098,938 | 14,838,431 | 92.2 |
| Bos taurus | GSE41637 | SRR594474 | Colon | 119,761,742 | 108,567,519 | 100,493,689 | 92.6 |
| | | SRR594475 | Heart | 117,554,231 | 102,808,834 | 99,087,699 | 96.4 |
| | | SRR594476 | Kidney | 115,720,336 | 107,048,435 | 102,311,713 | 95.6 |
| | | SRR594477 | Liver | 103,019,718 | 96,471,480 | 91,471,456 | 94.8 |
| | | SRR594478 | Lung | 103,196,444 | 94,356,596 | 89,631,315 | 95.0 |
| | | SRR594479 | Skeletal muscle | 113,227,169 | 100,258,839 | 95,034,372 | 94.8 |
| | | SRR594480 | Spleen | 116,884,523 | 106,487,738 | 98,708,719 | 92.7 |
| | | SRR594481 | Testes | 107,483,976 | 98,479,248 | 94,432,393 | 95.9 |
| | | SRR594483 | Colon | 32,336,017 | 26,359,467 | 23,192,961 | 88.0 |
| | | SRR594484 | Heart | 21,185,451 | 19,043,822 | 17,832,960 | 93.6 |
| | | SRR594485 | Kidney | 27,567,792 | 23,849,700 | 22,038,194 | 92.4 |
| | | SRR594486 | Liver | 26,021,524 | 22,615,391 | 20,906,823 | 92.5 |
| | | SRR594487 | Lung | 22,498,703 | 19,783,813 | 18,090,439 | 91.4 |
| | | SRR594488 | Skeletal muscle | 27,439,293 | 23,672,541 | 21,740,316 | 91.8 |
| | | SRR594489 | Spleen | 24,283,167 | 20,956,751 | 18,126,163 | 86.5 |
| | | SRR594490 | Testes | 38,368,885 | 28,286,737 | 25,950,324 | 91.7 |
| | | SRR594492 | Colon | 23,173,287 | 20,517,583 | 18,384,896 | 89.6 |
| | | SRR594493 | Heart | 37,897,106 | 29,975,705 | 28,038,599 | 93.5 |
| | | SRR594494 | Kidney | 22,535,945 | 19,780,690 | 18,307,870 | 92.6 |
| | | SRR594495 | Liver | 29,192,793 | 25,348,304 | 23,247,486 | 91.7 |
| | | SRR594496 | Lung | 35,817,736 | 28,434,425 | 26,123,049 | 91.9 |
| | | SRR594497 | Skeletal muscle | 24,842,489 | 21,711,317 | 20,211,951 | 93.1 |
| | | SRR594498 | Spleen | 31,925,721 | 25,398,752 | 21,828,414 | 85.9 |
| | | SRR594499 | Testes | 25,780,071 | 19,749,083 | 18,213,999 | 92.2 |
| Sus scrofa | ERP009821 | ERR789443 | Fat | 47,223,701 | 46,255,200 | 40,704,383 | 88.0 |
| | | ERR789444 | Heart | 46,474,210 | 45,751,115 | 40,401,277 | 88.3 |
| | | ERR789445 | Kidney | 47,565,661 | 46,737,485 | 41,432,264 | 88.7 |
| | | ERR789446 | Liver | 52,484,784 | 51,461,538 | 44,629,431 | 86.7 |
| | | ERR789447 | Lung | 42,115,278 | 41,036,112 | 33,759,440 | 82.3 |
| | | ERR789448 | Lymph node | 42,796,446 | 41,486,945 | 27,658,393 | 66.7 |
| | | ERR789449 | Muscle | 48,296,978 | 47,355,203 | 41,588,901 | 87.8 |
| | | ERR789450 | Spleen | 39,615,892 | 38,445,242 | 27,518,183 | 71.6 |
| Oryctolagus cuniculus | GSE106077 | SRR6206905 | Kidney | 59,053,890 | 50,965,385 | 44,586,791 | 87.5 |
| | | SRR6206925 | Testis | 64,474,890 | 55,281,359 | 47,267,601 | 85.5 |
| | | SRR6206910 | Liver | 55,990,721 | 48,037,240 | 42,156,774 | 87.8 |
| | | SRR6206900 | Heart | 55,064,609 | 47,251,883 | 40,998,926 | 86.8 |
| | | SRR6206920 | Skeletal muscle | 52,765,274 | 45,154,347 | 38,691,448 | 85.7 |
| | | SRR6206915 | Lung | 51,028,249 | 43,500,838 | 36,326,098 | 83.5 |
| Capra hircus | GSE43013 | SRR636844 | Liver | 15,873,867 | 15,539,416 | 14,327,927 | 92.2 |
| | | SRR636845 | Liver | 16,454,606 | 16,090,806 | 13,168,242 | 81.8 |
| | | SRR636894 | Kidney | 25,499,168 | 23,266,061 | 19,900,921 | 85.5 |
| | | SRR636895 | Kidney | 36,708,994 | 31,839,681 | 26,278,586 | 82.5 |

**Table S3. Alignment result of mammal ACE2 protein sequences in UniProt.**

```
A0A4X1UWQ8 PIG      MSGSFWLLLSLIPVTAAQSTTEELAKTFLEKFNLEAEDLAYQSSLASWNYNTNITDENIQ
K7GLM4 PIG          MSGSFWLLLSLIPVTAAQSTTEELAKTFLEKFNLEAEDLAYQSSLASWNYNTNITDENIQ
A0A452EVU0 CAPHI    MTGSFWLLLSLVAVTAAQSTTEEQAKTFLEKFNHEAEDLSYQSSLASWNYNTNITDENVQ
A0A4W2H6E0 BOBOX    MTGSFWLLLSLVAVTAAQSTTEEQAKTFLEKFNHEAEDLSYQSSLASWNYNTNITDENVQ
A0A4W2H3A1 BOBOX    MTGSFWLLLSLVAVTAAQSTTEEQAKTFLEKFNHEAEDLSYQSSLASWNYNTNITDENVQ
Q2HJI5 BOVIN        MTGSFWLLLSLVAVTAAQSTTEEQAKTFLEKFNHEAEDLSYQSSLASWNYNTNITDENVQ
ACE2 BOVIN          MTGSFWLLLSLVAVTAAQSTTEEQAKTFLEKFNHEAEDLSYQSSLASWNYNTNITDENVQ
A0A452EVJ5 CAPHI    MTGSFWLLLSLVAVTAAQSTTEEQAKTFLEKFNHEAEDLSYQSSLASWNYNTNITDENVQ
W5PSB6 SHEEP        MTGSFWLLLSLVAVTAAQSTTEGQAKTFLEKFNHEAEDLSYQSSLASWNYNTNITDENVQ
E2DHI7 9CHIR        MSGSFWLLLSLVAVTTAQSTTEDRAKTFLDEFNSEAENLSYQSSLASWDYNTNINDENVQ
E2DHI4 9CHIR        MSGSFWFLLSLVAVTTAQSTTEDEAKIFLDKFNTKAEDLSHQSSLASWDYNTNINDENVQ
U5WHY8 9CHIR        MSGSSWLLLSLVAVTTAQSTTEDEAKMFLDKFNTKAEDLSHQSSLASWDYNTNINDENVQ
G1TEF4 RABIT        MSGSSWLLLSLVAVTAAQSTIEELAKTFLEKFNQEAEDLSYQSALASWDYNTNITEENVQ
A0A1U7QTA1 MESAU    MSSSSWLLLSLVAVTTAQSIIEEQAKTFLDKFNQEAEDLSYQSALASWNYNTNITEENAQ
ACE2 MOUSE          MSSSSWLLLSLVAVTTAQSLTEENAKTFLNNFNQEAEDLSYQSSLASWNYNTNITEENAQ
ACE2 RAT            MSSSCWLLLSLVAVATAQSLIEEKAESFLNKFNQEAEDLSYQSSLASWNYNTNITEENAQ
ACE2 HUMAN          MSSSSWLLLSLVAVTAAQSTIEEQAKTFLDKFNHEAEDLFYQSSLASWNYNTNITEENVQ
A0A384DV19 FELCA    MSGSFWLLLSFAALTAAQSTTEELAKTFLEKFNHEAEELSYQSSLASWNYNTNITDENVQ
A0A5F5XDN9 FELCA    MSGSFWLLLSFAALTAAQSTTEELAKTFLEKFNHEAEELSYQSSLASWNYNTNITDENVQ
ACE2 FELCA          MSGSFWLLLSFAALTAAQSTTEELAKTFLEKFNHEAEELSYQSSLASWNYNTNITDENVQ
A0A5F4CXG9 CANLF    MSGSSWLLLSLAALTAAQS-TEDLVKTFLEKFNYEAEELSYQSSLASWNYNINITDENVQ
A0A5F4BS93 CANLF    MSGSSWLLLSLAALTAAQS-TEDLVKTFLEKFNYEAEELSYQSSLASWNYNINITDENVQ
F1P7C5 CANLF        MSGSSWLLLSLAALTAAQS-TEDLVKTFLEKFNYEAEELSYQSSLASWNYNINITDENVQ
J9P7Y2 CANLF        MSGSSWLLLSLAALTAAQS-TEDLVKTFLEKFNYEAEELSYQSSLASWNYNINITDENVQ
                    *:.* *:***:  :::***  *  .: **::** :**:* :**:****:** **.:** *

A0A4X1UWQ8 PIG      KMNDARAKWSAFYEEQSRIAKTYPLDEIQTLILKRQLQALQQSGTSGLSADKSKRLNTIL
K7GLM4 PIG          KMNDARAKWSAFYEEQSRIAKTYPLDEIQTLILKRQLQALQQSGTSGLSADKSKRLNTIL
A0A452EVU0 CAPHI    KMNEARAKWSAFYEEQSRMARTYSLEEIQNLTLKRQLKALQHSGTSVLSAEKSKRVYFIE
A0A4W2H6E0 BOBOX    KMNEARAKWSAFYEEQSRMAKTYSLEEIQNLTLKRQLKALQHSGTSALSAEKSKRLNTIL
A0A4W2H3A1 BOBOX    KMNEARAKWSAFYEEQSRMAKTYSLEEIQNLTLKRQLKALQHSGTSALSAEKSKRLNTIL
Q2HJI5 BOVIN        KMNEARAKWSAFYEEQSRMAKTYSLEEIQNLTLKRQLKALQHSGTSALSAEKSKRLNTIL
ACE2 BOVIN          KMNEARAKWSAFYEEQSRMAKTYSLEEIQNLTLKRQLKALQHSGTSALSAEKSKRLNTIL
A0A452EVJ5 CAPHI    KMNEARAKWSAFYEEQSRMARTYSLEEIQNLTLKRQLKALQHSGTSVLSAEKSKRLNMIL
W5PSB6 SHEEP        KMNEARAKWSAFYEEQSRMARTYSLEEIQNLTLKRQLKALQHSGTSVLSAEKSKRLNTIL
E2DHI7 9CHIR        KMDEAGAKWSAFYEEQSKLAKNYPLEQIQNVTVKLQLQILQQSGSPVLSEDKSKRLNSIL
E2DHI4 9CHIR        KMDEAGAKWSAFYEEQSKLAKNYPLEQIQNVTVKLQLQILQQSGSPVLSEDKSKRLNSIL
U5WHY8 9CHIR        KMDEAGAKWSAFYEEQSKLAKNYSLEQIQNVTVKLQLQILQQSGSPVLSEDKSKRLNSIL
G1TEF4 RABIT        KMNDAEAKWSAFYEEQSKLAKTYPSQEVQNLTVKRQLQALQQSGSSALSADKSKQLNTIL
A0A1U7QTA1 MESAU    KMNEAAAKWSAFYEEQSKLAKNYSLQEVQNLTIKRQLQALQQSGSSALSADKNKQLNTIL
ACE2 MOUSE          KMSEAAAKWSAFYEEQSKTAQSFSLQEIQTPIIKRQLQALQQSGSSALSADKNKQLNTIL
ACE2 RAT            KMNEAAAKWSAFYEEQSKIAQNFSLQEIQNATIKRQLKALQQSGSSALSPDKNKQLNTIL
ACE2 HUMAN          NMNNAGDKWSAFLKEQSTLAQMYPLQEIQNLTVKLQLQALQQNGSSVLSEDKSKRLNTIL
A0A384DV19 FELCA    KMNEAGAKWSAFYEEQSKLAKTYPLAEIHNTTVKRQLQALQQSGSSVLSADKSQRLNTIL
A0A5F5XDN9 FELCA    KMNEAGAKWSAFYEEQSKLAKTYPLAEIHNTTVKRQLQALQQSGSSVLSADKSQRLNTIL
ACE2 FELCA          KMNEAGAKWSAFYEEQSKLAKTYPLAEIHNTTVKRQLQALQQSGSSVLSADKSQRLNTIL
A0A5F4CXG9 CANLF    KMNNAGAKWSAFYEEQSKLAKTYPLEEIQDSTVKRQLRALQHSGSSVLSADKNQRLNTIL
A0A5F4BS93 CANLF    KMNNAGAKWSAFYEEQSKLAKTYPLEEIQDSTVKRQLRALQHSGSSVLSADKNQRLNTIL
F1P7C5 CANLF        KMNNAGAKWSAFYEEQSKLAKTYPLEEIQDSTVKRQLRALQHSGSSVLSADKNQRLNTIL
J9P7Y2 CANLF        KMNNAGAKWSAFYEEQSKLAKTYPLEEIQDSTVKRQLRALQHSGSSVLSADKNQRLNTIL
                    :*.:*  ***** :***  *: :   :::   :* **: **:.*:  ** :*.:::  *

A0A4X1UWQ8 PIG      NTMSTIYSSGKVLDPNNPQECLVLEPGLDEIMENSKDYSRRLWAWESWRAEVGKQLRPLY
K7GLM4 PIG          NTMSTIYSSGKVLDPNNPQECLVLEPGLDEIMENSKDYSRRLWAWESWRAEVGKQLRPLY
A0A452EVU0 CAPHI    Q-------------------LYDLDICLDDIMENSRDYNRRLWAWEGWRAEVGKQLRPLY
A0A4W2H6E0 BOBOX    NKMSTIYSTGKVLDPN-TQECLALEPGLDDIMENSRDYNRRLWAWEGWRAEVGKQLRPLY
A0A4W2H3A1 BOBOX    NKMSTIYSTGKVLDPN-TQECLALEPGLDDIMENSRDYNRRLWAWEGWRAEVGKQLRPLY
Q2HJI5 BOVIN        NKMSTIYSTGKVLDPN-TQECLALEPGLDDIMENSRDYNRRLWAWEGWRAEVGKQLRPLY
```

```
ACE2 BOVIN         NKMSTIYSTGKVLDPN-TQECLALEPGLDDIMENSRDYNRRLWAWEGWRAEVGKQLRPLY
A0A452EVJ5 CAPHI   NKMSTIYSTGKVLDPN-TQECLALEPGLDDIMENSRDYNRRLWAWEGWRAEVGKQLRPLY
W5PSB6 SHEEP       NKMSTIYSTGKVLDPN-TQECLALEPGLDDIMENSRDYNRRLWAWEGWRAEVGKQLRPLY
E2DHI7 9CHIR       NAMSTIYSTGKVCKPNKPHECLLLEPGLDNIMGTSKDYNERLWAWEGWRAEVGKQLRPLY
E2DHI4 9CHIR       NAMSTIYSTGKVCKPNKPHECLLLEPGLDNIMGTSKDYNERLWAWEGWRAEVGKQLRPLY
U5WHY8 9CHIR       NAMSTIYSTGKVCKPNKPQECLLLEPGLDNIMGTSKDYNERLWAWEGWRAEVGKQLRPLY
G1TEF4 RABIT       STMSTIYSTGKVCNQSNPQECFLLEPGLDEIMAKSTDYNERLWAWEGWRSVVGKQLRPLY
A0A1U7QTA1 MESAU   NTMSTIYSTGKVCNPKNPQECLLLEPGLDDIMATSTDYNERLWAWEGWRAEVGKQLRPLY
ACE2 MOUSE         NTMSTIYSTGKVCNPKNPQECLLLEPGLDEIMATSTDYNSRLWAWEGWRAEVGKQLRPLY
ACE2 RAT           NTMSTIYSTGKVCNSMNPQECFLLEPGLDEIMATSTDYNRRLWAWEGWRAEVGKQLRPLY
ACE2 HUMAN         NTMSTIYSTGKVCNPDNPQECLLLEPGLNEIMANSLDYNERLWAWESWRSEVGKQLRPLY
A0A384DV19 FELCA   NAMSTIYSTGKACNPNNPQECLLLEPGLDDIMENSKDYNERLWAWEGWRAEVGKQLRPLY
A0A5F5XDN9 FELCA   NAMSTIYSTGKACNPNNPQECLLLEPGLDDIMENSKDYNERLWAWEGWRAEVGKQLRPLY
ACE2 FELCA         NAMSTIYSTGKACNPNNPQECLLLEPGLDDIMENSKDYNERLWAWEGWRAEVGKQLRPLY
A0A5F4CXG9 CANLF   NSMSTIYSTGKACNPSNPQECLLLEPGLDDIMENSKDYNERLWAWEGWRSEVGKQLRPLY
A0A5F4BS93 CANLF   NSMSTIYSTGKACNPSNPQECLLLEPGLDDIMENSKDYNERLWAWEGWRSEVGKQLRPLY
F1P7C5 CANLF       NSMSTIYSTGKACNPSNPQECLLLEPGLDDIMENSKDYNERLWAWEGWRSEVGKQLRPLY
J9P7Y2 CANLF       NSMSTIYSTGKACNPSNPQECLLLEPGLDDIMENSKDYNERLWAWEGWRSEVGKQLRPLY
                   .                  *:  *::** .* **. ******.**: *********

A0A4X1UWQ8 PIG     EEYVVLENEMARAN-----------NYEDYGDYWRGDYEVTGTGDYDYSRNQLMEDVERT
K7GLM4 PIG         EEYVVLENEMARAN-----------NYEDYGDYWRGDYEVTGTGDYDYSRNQLMEDVERT
A0A452EVU0 CAPHI   EEYVVLENEMARANSKCSHFLLFETDYEDYGDYWRGDYEVTGAGDYDYSRDQLMKDVERT
A0A4W2H6E0 BOBOX   EEYVVLENEMARAN-----------NYEDYGDYWRGDYEVTGAGDYDYSRDQLMKDVERT
A0A4W2H3A1 BOBOX   EEYVVLENEMARAN-----------NYEDYGDYWRGDYEVTGAGDYDYSRDQLMKDVERT
Q2HJI5 BOVIN       EEYVVLENEMARAN-----------NYEDYGDYWRGDYEVTGAGDYDYSRDQLMKDVERT
ACE2 BOVIN         EEYVVLENEMARAN-----------NYEDYGDYWRGDYEVTGAGDYDYSRDQLMKDVERT
A0A452EVJ5 CAPHI   EEYVVLENEMARAN-----------NYEDYGDYWRGDYEVTGAGDYDYSRDQLMKDVERT
W5PSB6 SHEEP       EEYVVLENEMARAN-----------NYEDYGDYWRGDYEVTGAGDYDYSRDQLMKDVERT
E2DHI7 9CHIR       EEYVVLKNEMARGY-----------HYEDYGDYWRRDYETEESPGPGYSRDQLMKDVERI
E2DHI4 9CHIR       EEYVVLKNEMARGY-----------HYEDYGDYWRRDYETEESPGPGYSRDQLMKDVERI
U5WHY8 9CHIR       EEYVVLKNEMARGY-----------HYEDYGDYWRRDYETEESPGPGYSRDQLMKDVERI
G1TEF4 RABIT       EEYVVLKNEMARAN-----------NYEDYGDYWRADYEAEGADGYDYSRSQLIDDVERT
A0A1U7QTA1 MESAU   EEYVVLKNEMARAN-----------NYEDYGDYWRGDYEAEGADGYNYNGNQLIEDVERT
ACE2 MOUSE         EEYVVLKNEMARAN-----------NYNDYGDYWRGDYEAEGADGYNYNRNQLIEDVERT
ACE2 RAT           EEYVVLKNEMARAN-----------NYEDYGDYWRGDYEAEGVEGYNYNRNQLIEDVENT
ACE2 HUMAN         EEYVVLKNEMARAN-----------HYEDYGDYWRGDYEVNGVDGYDYSRGQLIEDVEHT
A0A384DV19 FELCA   EEYVALKNEMAK-------------NYEDYGDYWRGDYEEEWTDGYNYSRSQLIKDVEHT
A0A5F5XDN9 FELCA   EEYVALKNEMAKSK-----------QYEDYGDYWRGDYEEEWTDGYNYSRSQLIKDVEHT
ACE2 FELCA         EEYVALKNEMARAN-----------NYEDYGDYWRGDYEEEWTDGYNYSRSQLIKDVEHT
A0A5F4CXG9 CANLF   EEYVALKNEMARAN-----------NYEDYGDYWRGDYEEEWENGYNYSRNQLIDDVEHT
A0A5F4BS93 CANLF   EEYVALKNEMARAN-----------NYEDYGDYWRGDYEEEWENGYNYSRNQLIDDVEHT
F1P7C5 CANLF       EEYVALKNEMARAN-----------NYEDYGDYWRGDYEEEWENGYNYSRNQLIDDVEHT
J9P7Y2 CANLF       EEYVALKNEMARAN-----------NYEDYGDYWRGDYEEEWENGYNYSRNQLIDDVEHT
                   ****.*:****:           .*:******* ***    . .*. .**:.***.

A0A4X1UWQ8 PIG     FAEIKPLYEHLHAYVRAKLMDAYPSRISPTGCLPAHL-LGDMWGRFWTNLYPLTVPFGEK
K7GLM4 PIG         FAEIKPLYEHLHAYVRAKLMDAYPSRISPTGCLPAHL-LGDMWGRFWTNLYPLTVPFGEK
A0A452EVU0 CAPHI   FAEIKPLYEQLHAYVRAKLMNTYPSYISPTGCLPAHL-LGDMWGRFWTNLYSLTVPFEHK
A0A4W2H6E0 BOBOX   FAEIKPLYEQLHAYVRAKLMHTYPSYISPTGCLPAHL-LGDMWGRFWTNLYSLTVPFEHK
A0A4W2H3A1 BOBOX   FAEIKPLYEQLHAYVRAKLMHTYPSYISPTGCLPAHL-LGDMWGRFWTNLYSLTVPFEHK
Q2HJI5 BOVIN       FAEIKPLYEQLHAYVRAKLMHTYPSYISPTGCLPAHL-LGDMWGRFWTNLYSLTVPFEHK
ACE2 BOVIN         FAEIKPLYEQLHAYVRAKLMHTYPSYISPTGCLPAHL-LGDMWGRFWTNLYSLTVPFEHK
A0A452EVJ5 CAPHI   FAEIKPLYEQLHAYVRAKLMNTYPSYISPTGCLPAHL-LGDMWGRFWTNLYSLTVPFEHK
W5PSB6 SHEEP       FEEIKPLYEQLHAYVRAKLMDTYPSYISPTGCLPAHLLVGDMWGRFWTNLYSLTVPFEHK
E2DHI7 9CHIR       FTEIKPLYEHLHAYVRAKLMDTYPFHISPTGCLPAHL-LGDMWGRFWTNLYPLTVPFGQK
E2DHI4 9CHIR       FTEIKPLYEHLHAYVRAKLMDTYPFHISPTGCLPAHL-LGDMWGRFWTNLYPLTVPFGQK
U5WHY8 9CHIR       FTEIKPLYEHLHAYVRAKLMDTYPFHISPTGCLPAHL-LGDMWGRFWTNLYPLTVPFGQK
G1TEF4 RABIT       FSEIKPLYEQLHAFVRTKLMDAYPSRISPTGCLPAHL-LGDMWGRFWTNLYSLTVPFGQK
A0A1U7QTA1 MESAU   FKEIKPLYEQLHAYVRTKLMNTYPSYISPTGCLPAHL-LGDMWGRFWTNLYPLTVPFGQK
ACE2 MOUSE         FAEIKPLYEHLHAYVRRKLMDTYPSYISPTGCLPAHL-LGDMWGRFWTNLYPLTVPFAQK
```

```
ACE2 RAT          FKEIKPLYEQLHAYVRTKLMEVYPSYISPTGCLPAHL-LGDMWGRFWTNLYPLTTPFLQK
ACE2 HUMAN        FEEIKPLYEHLHAYVRAKLMNAYPSYISPIGCLPAHL-LGDMWGRFWTNLYSLTVPFGQK
A0A384DV19 FELCA  FTQIKPLYQHLHAYVRAKLMDTYPSRISPTGCLPAHL-LGDMWGRFWTNLYPLTVPFGQK
A0A5F5XDN9 FELCA  FTQIKPLYQHLHAYVRAKLMDTYPSRISPTGCLPAHL-LGDMWGRFWTNLYPLTVPFGQK
ACE2 FELCA        FTQIKPLYQHLHAYVRAKLMDTYPSRISPTGCLPAHL-LGDMWGRFWTNLYPLTVPFGQK
A0A5F4CXG9 CANLF  FTQIMPLYQHLHAYVRTKLMDTYPSYISPTGCLPAHL-LGDMWGRFWTNLYPLTVPFGQK
A0A5F4BS93 CANLF  FTQIMPLYQHLHAYVRTKLMDTYPSYISPTGCLPAHL-LGDMWGRFWTNLYPLTVPFGQK
F1P7C5 CANLF      FTQIMPLYQHLHAYVRTKLMDTYPSYISPTGCLPAHL-LGDMWGRFWTNLYPLTVPFGQK
J9P7Y2 CANLF      FTQIMPLYQHLHAYVRTKLMDTYPSYISPTGCLPAHL-LGDMWGRFWTNLYPLTVPFGQK
                  * :* ***::***:** ***..**  *** ******* :************ **.** .*

A0A4X1UWQ8 PIG    PSIDVTEAMVNQSWDAVRIFEEAEKFFVSIGLPNMTQGFWNNSMLTEPGDGRKVVCHPTA
K7GLM4 PIG        PSIDVTEAMVNQSWDAIRIFEEAEKFFVSIGLPNMTQGFWNNSMLTEPGDGRKVVCHPTA
A0A452EVU0 CAPHI  PSIDVTEKMKNQSWDAERIFKEAEKFFVSIGLPYMTQGFWNNSMLTEPGDGRKVVCHPTA
A0A4W2H6E0 BOBOX  PSIDVTEKMENQSWDAERIFKEAEKFFVSISLPYMTQGFWDNSMLTEPGDGRKVVCHPTA
A0A4W2H3A1 BOBOX  PSIDVTEKMENQSWDAERIFKEAEKFFVSISLPYMTQGFWDNSMLTEPGDGRKVVCHPTA
Q2HJI5 BOVIN      PSIDVTEKMENQSWDAERIFKEAEKFFVSISLPYMTQGFWDNSMLTEPGDGRKVVCHPTA
ACE2 BOVIN        PSIDVTEKMENQSWDAERIFKEAEKFFVSISLPYMTQGFWDNSMLTEPGDGRKVVCHPTA
A0A452EVJ5 CAPHI  PSIDVTEKMKNQSWDAERIFKEAEKFFVSIGLPYMTQGFWNNSMLTEPGDGRKVVCHPTA
W5PSB6 SHEEP      PSIDVTEKMKNQSWDAERIFKEAEKFFVSIGLPYMTQGFWDNSMLTEPGDGRKVVCHPTA
E2DHI7 9CHIR      PNIDVTDEMLKQGWDADRIFKEAEKFFVSVGLPNMTEGFWNNSMLTEPGDGRKVVCHPTA
E2DHI4 9CHIR      PNIDVTDEMLKQGWDADRIFKEAEKFFVSVGLPNMTEGFWNNSMLTEPGDGRKVVCHPTA
U5WHY8 9CHIR      PNIDVTDEMLKQGWDADRIFKEAEKFFVSVGLPNMTEGFWNNSMLTEPGDGRKVVCHPTA
G1TEF4 RABIT      PNIDVTDTMVNQGWDAERIFKEAEKFFVSVGLPSMTQGFWENSMLTEPGDGRKVVCHPTA
A0A1U7QTA1 MESAU  PNIDVTDAMVNQGWNAERIFKEAEKFFVSVGLPYMTQGFWENSMLTDPGDDRKVVCHPTA
ACE2 MOUSE        PNIDVTDAMMNQGWDAERIFQEAEKFFVSVGLPHMTQGFWANSMLTEPADGRKVVCHPTA
ACE2 RAT          PNIDVTDAMVNQSWDAERIFKEAEKFFVSVGLPQMTPGFWTNSMLTEPGDDRKVVCHPTA
ACE2 HUMAN        PNIDVTDAMVDQAWDAQRIFKEAEKFFVSVGLPNMTQGFWENSMLTDPGNVQKAVCHPTA
A0A384DV19 FELCA  PNIDVTDAMVNQSWDARRIFKEAEKFFVSVGLPNMTQGFWENSMLTEPGDSRKVVCHPTA
A0A5F5XDN9 FELCA  PNIDVTDAMVNQSWDARRIFKEAEKFFVSVGLPNMTQGFWENSMLTEPGDSRKVVCHPTA
ACE2 FELCA        PNIDVTDAMVNQSWDARRIFKEAEKFFVSVGLPNMTQGFWENSMLTEPGDSRKVVCHPTA
A0A5F4CXG9 CANLF  PNIDVTNAMVNQSWDARKIFKEAEKFFVSVGLPNMTQEFWENSMLTEPSDSRKVVCHPTA
A0A5F4BS93 CANLF  PNIDVTNAMVNQSWDARKIFKEAEKFFVSVGLPNMTQEFWENSMLTEPSDSRKVVCHPTA
F1P7C5 CANLF      PNIDVTNAMVNQSWDARKIFKEAEKFFVSVGLPNMTQEFWENSMLTEPSDSRKVVCHPTA
J9P7Y2 CANLF      PNIDVTNAMVNQSWDARKIFKEAEKFFVSVGLPNMTQEFWENSMLTEPSDSRKVVCHPTA
                  *.****: * .*.*:* :**:********:.** **  ** *****:*.: :*.******

A0A4X1UWQ8 PIG    WDLGKGDFRIKMCTKVTMDDFLTAHHEMGHIQYDMAYAIQPYLLRNGANEGFHEAVGEIM
K7GLM4 PIG        WDLGKGDFRIKMCTKVTMDDFLTAHHEMGHIQYDMAYAIQPYLLRNGANEGFHEAVGEIM
A0A452EVU0 CAPHI  WDLGKGDFRIKMCTKVTMDDFLTAHHEMGHIQYDMAYATQPYLLRNGANEGFHEAVGEIM
A0A4W2H6E0 BOBOX  WDLGKGDFRIKMCTKVTMDDFLTAHHEMGHIQYDMAYAAQPYLLRNGANEGFHEAVGEIM
A0A4W2H3A1 BOBOX  WDLGKGDFRIKMCTKVTMDDFLTAHHEMGHIQYDMAYAAQPYLLRNGANEGFHEAVGEIM
Q2HJI5 BOVIN      WDLGKGDFRIKMCTKVTMDDFLTAHHEMGHIQYDMAYAAQPYLLRNGANEGFHEAVGEIM
ACE2 BOVIN        WDLGKGDFRIKMCTKVTMDDFLTAHHEMGHIQYDMAYAAQPYLLRNGANEGFHEAVGEIM
A0A452EVJ5 CAPHI  WDLGKGDFRIKMCTKVTMDDFLTAHHEMGHIQYDMAYATQPYLLRNGANEGFHEAVGEIM
W5PSB6 SHEEP      WDLGKGDFRIKMCTKVTMDDFLTAHHEMGHIQYDMAYATQPYLLRNGANEGFHEAVGEIM
E2DHI7 9CHIR      WDLGKGDFRIKMCTKVTMEDFLTAHHEMGHIQYDMVYASQPYLLRNGANEGFHEAVGEVM
E2DHI4 9CHIR      WDLGKGDFRIKMCTKVTMEDFLTAHHEMGHIQYDMAYASQPYLLRNGANEGFHEAVGEVM
U5WHY8 9CHIR      WDLGKGDFRIKMCTKVTMEDFLTAHHEMGHIQYDMAYASQPYLLRNGANEGFHEAVGEVM
G1TEF4 RABIT      WDLGKGDFRIKMCTKVTMDNFLTAHHEMGHIQYDMAYATQPFLLRNGANEGFHEAVGEIM
A0A1U7QTA1 MESAU  WDLGKGDFRIKMCTKVTMDNFLTAHHEMGHIQYDMAYATQPFLLRNGANEGFHEAVGEIM
ACE2 MOUSE        WDLGHGDFRIKMCTKVTMDNFLTAHHEMGHIQYDMAYARQPFLLRNGANEGFHEAVGEIM
ACE2 RAT          WDLGHGDFRIKMCTKVTMDNFLTAHHEMGHIQYDMAYAKQPFLLRNGANEGFHEAVGEIM
ACE2 HUMAN        WDLGKGDFRILMCTKVTMDDFLTAHHEMGHIQYDMAYAAQPFLLRNGANEGFHEAVGEIM
A0A384DV19 FELCA  WDLGKGDFRIKMCTKVTMDDFLTAHHEMGHIQYDMAYAVQPFLLRNGANEGFHEAVGEIM
A0A5F5XDN9 FELCA  WDLGKGDFRIKMCTKVTMDDFLTAHHEMGHIQYDMAYAVQPFLLRNGANEGFHEAVGEIM
ACE2 FELCA        WDLGKGDFRIKMCTKVTMDDFLTAHHEMGHIQYDMAYAVQPFLLRNGANEGFHEAVGEIM
A0A5F4CXG9 CANLF  WDLGKGDFRIKMCTKVTMDDFLTAHHEMGHIQYDMAYAAQPFLLRNGANEGFHEAVGEIM
A0A5F4BS93 CANLF  WDLGKGDFRIKMCTKVTMDDFLTAHHEMGHIQYDMAYAAQPFLLRNGANEGFHEAVGEIM
F1P7C5 CANLF      WDLGKGDFRIKMCTKVTMDDFLTAHHEMGHIQYDMAYAAQPFLLRNGANEGFHEAVGEIM
J9P7Y2 CANLF      WDLGKGDFRIKMCTKVTMDDFLTAHHEMGHIQYDMAYAAQPFLLRNGANEGFHEAVGEIM
```

```
                    ****:***** *******::***************.** **:****************:*

A0A4X1UWQ8 PIG      SLSAATPHYLKALGLLPPDFYEDSETEINFLLKQALTIVGTLPFTYMLEKWRWMVFKGEI
K7GLM4 PIG          SLSAATPHYLKALGLLPPDFYEDSETEINFLLKQALTIVGTLPFTYMLEKWRWMVFKGEI
A0A452EVU0 CAPHI    SLSAATPHYLKALGLLAPDFYEDNETEINFLLKQALTIVGTLPFTYMLEKWRWMVFKGEI
A0A4W2H6E0 BOBOX    SLSAATPHYLKALGLLAPDFHEDNETEINFLLKQALTIVGTLPFTYMLEKWRWMVFKGEI
A0A4W2H3A1 BOBOX    SLSAATPHYLKALGLLAPDFHEDNETEINFLLKQALTIVGTLPFTYMLEKWRWMVFKGEI
Q2HJI5 BOVIN        SLSAATPHYLKALGLLAPDFHEDNETEINFLLKQALTIVGTLPFTYMLEKWRWMVFKGEI
ACE2 BOVIN          SLSAATPHYLKALGLLAPDFHEDNETEINFLLKQALTIVGTLPFTYMLEKWRWMVFKGEI
A0A452EVJ5 CAPHI    SLSAATPHYLKALGLLAPDFYEDNETEINFLLKQALTIVGTLPFTYMLEKWRWMVFKGEI
W5PSB6 SHEEP        SLSAATPHYLKALGLLAPDFYEDNETEINFLLKQALTIVGTLPFTYMLEKWRWMVFKGEI
E2DHI7 9CHIR        SLSVATPKHLKTMGLLSPDFHEDNETEINFLLKQALNIVGTLPFTYMLEKWRWMVFKGEI
E2DHI4 9CHIR        SLSVATPKHLKTMGLLSPDFREDNETEINFLLKQALNIVGTLPFTYMLEKWMWMVFKGEI
U5WHY8 9CHIR        SLSVATPKHLKTMGLLSPDFREDNETEINFLLKQALNIVGTLPFTYMLEKWRWMVFKGEI
G1TEF4 RABIT        SLSAATPEHLKSIGLLPYDFHEDNETEINFLLKQALTIVGTLPFTYMLEKWRWMVFKGEI
A0A1U7QTA1 MESAU    SLSAATPEHLKSIGLLPSDFQEDNETEINFLLKQALTIVGTLPFTYMLEKWRWMVFKGDI
ACE2 MOUSE          SLSAATPKHLKSIGLLPSDFQEDSETEINFLLKQALTIVGTLPFTYMLEKWRWMVFRGEI
ACE2 RAT            SLSAATPKHLKSIGLLPSNFQEDNETEINFLLKQALTIVGTLPFTYMLEKWRWMVFQDKI
ACE2 HUMAN          SLSAATPKHLKSIGLLSPDFQEDNETEINFLLKQALTIVGTLPFTYMLEKWRWMVFKGEI
A0A384DV19 FELCA    SLSAATPNHLKTIGLLSPGFSEDSETEINFLLKQALTIVGTLPFTYMLEKWRWMVFKGEI
A0A5F5XDN9 FELCA    SLSAATPNHLKTIGLLSPGFSEDSETEINFLLKQALTIVGTLPFTYMLEKWRWMVFKGEI
ACE2 FELCA          SLSAATPNHLKTIGLLSPGFSEDSETEINFLLKQALTIVGTLPFTYMLEKWRWMVFKGEI
A0A5F4CXG9 CANLF    SLSAATPNHLKNIGLLPPSFFEDSETEINFLLKQALTIVGTLPFTYMLEKWRWMVFKGEI
A0A5F4BS93 CANLF    SLSAATPNHLKNIGLLPPSFFEDSETEINFLLKQALTIVGTLPFTYMLEKWRWMVFKGEI
F1P7C5 CANLF        SLSAATPNHLKNIGLLPPSFFEDSETEINFLLKQALTIVGTLPFTYMLEKWRWMVFKGEI
J9P7Y2 CANLF        SLSAATPNHLKNIGLLPPSFFEDSETEINFLLKQALTIVGTLPFTYMLEKWRWMVFKGEI
                    ***.***.:** :***  .* **.************.************** ****:..*

A0A4X1UWQ8 PIG      PKEQWMQKWWEMKREIVGVVEPLPHDETYCDPACLFHVAEDYSFIRYYTRTIYQFQFHEA
K7GLM4 PIG          PKEQWMQKWWEMKREIVGVVEPLPHDETYCDPACLFHVAEDYSFIRYYTRTIYQFQFHEA
A0A452EVU0 CAPHI    PKQQWMEKWWEMKREIVGVVEPLPHDETYCDPACLFHVAEDYSFIRYYTRTIYQFQFHEA
A0A4W2H6E0 BOBOX    PKQQWMEKWWEMKREIVGVVEPLPHDETYCDPACLFHVAEDYSFIRYYTRTIYQFQFHEA
A0A4W2H3A1 BOBOX    PKQQWMEKWWEMKREIVGVVEPLPHDETYCDPACLFHVAEDYSFIRYYTRTIYQFQFHEA
Q2HJI5 BOVIN        PKQQWMEKWWEMKREIVGVVEPLPHDETYCDPACLFHVAEDYSFIRYYTRTIYQFQFHEA
ACE2 BOVIN          PKQQWMEKWWEMKREIVGVVEPLPHDETYCDPACLFHVAEDYSFIRYYTRTIYQFQFHEA
A0A452EVJ5 CAPHI    PKQQWMEKWWEMKREIVGVVEPLPHDETYCDPACLFHVAEDYSFIRYYTRTIYQFQFHEA
W5PSB6 SHEEP        PKQQWMEKWWEMKREIVGVVEPLPHDETYCDPACLFHVAEDYSFIRYYTRTIYQFQFHEA
E2DHI7 9CHIR        PKEEWMKKWWEMKRKIVGVVEPVPHDETYCDPASLFHVANDYSFIRYYTRTIFEFQFHEA
E2DHI4 9CHIR        PKEEWMKKWLEMKRKIVGVVEPVPHDETYCDPASLFHVANDYSFIRYYTRTIFEFQFHEA
U5WHY8 9CHIR        PKEEWMKKWWEMKRKIVGVVEPVPHDETYCDPASLFHVANDYSFIRYYTRTIFEFQFHEA
G1TEF4 RABIT        PKEQWMQKWWEMKREIVGVVEPMPHDETYCDPAALFHVANDYSFIRYYTRTIYQFQFQEA
A0A1U7QTA1 MESAU    PKEQWMEKWWEMKREIVGVVEPLPHDETYCDPAALFHVSNDYSFIRYYTRTIYQFQFQEA
ACE2 MOUSE          PKEQWMKKWWEMKREIVGVVEPLPHDETYCDPASLFHVSNDYSFIRYYTRTIYQFQFQEA
ACE2 RAT            PREQWTKKWWEMKREIVGVVEPLPHDETYCDPASLFHVSNDYSFIRYYTRTIYQFQFQEA
ACE2 HUMAN          PKDQWMKKWWEMKREIVGVVEPVPHDETYCDPASLFHVSNDYSFIRYYTRTLYQFQFQEA
A0A384DV19 FELCA    PKEQWMQKWWEMKREIVGVVEPVPHDETYCDPASLFHVANDYSFIRYYTRTIYQFQFQEA
A0A5F5XDN9 FELCA    PKEQWMQKWWEMKREIVGVVEPVPHDETYCDPASLFHVANDYSFIRYYTRTIYQFQFQEA
ACE2 FELCA          PKEQWMQKWWEMKREIVGVVEPVPHDETYCDPASLFHVANDYSFIRYYTRTIYQFQFQEA
A0A5F4CXG9 CANLF    PKDQWMKTWWEMKRNIVGVVEPVPHDETYCDPASLFHVANDYSFIRYYTRTIYQFQFQEA
A0A5F4BS93 CANLF    PKDQWMKTWWEMKRNIVGVVEPVPHDETYCDPASLFHVANDYSFIRYYTRTIYQFQFQEA
F1P7C5 CANLF        PKDQWMKTWWEMKRNIVGVVEPVPHDETYCDPASLFHVANDYSFIRYYTRTIYQFQFQEA
J9P7Y2 CANLF        PKDQWMKTWWEMKRNIVGVVEPVPHDETYCDPASLFHVANDYSFIRYYTRTIYQFQFQEA
                    *:::* :.* ****:*******:**********.****::***********:::***:**

A0A4X1UWQ8 PIG      LCRTAKHEGPLYKCDISNSTEAGQKLLQMLSLGKSEPWTLALENIVGVKTMDVKPLLSYF
K7GLM4 PIG          LCRTAKHEGPLYKCDISNSTEAGQKLLQMLSLGKSEPWTLALENIVGVKTMDVKPLLSYF
A0A452EVU0 CAPHI    LCKTAKHEGALFKCDISNSTEAGQRLLQMLRLGKSEPWTLALENIVGIKTMDVKPLLNYF
A0A4W2H6E0 BOBOX    LCKTAKHEGALFKCDISNSTEAGQRLLQMLRLGKSEPWTLALENIVGIKTMDVKPLLNYF
A0A4W2H3A1 BOBOX    LCKTAKHEGALFKCDISNSTEAGQRLLQMLRLGKSEPWTLALENIVGIKTMDVKPLLNYF
Q2HJI5 BOVIN        LCKTAKHEGALFKCDISNSTEAGQRLLQMLRLGKSEPWTLALENIVGIKTMDVKPLLNYF
ACE2 BOVIN          LCKTAKHEGALFKCDISNSTEAGQRLLQMLRLGKSEPWTLALENIVGIKTMDVKPLLNYF
```

```
A0A452EVJ5 CAPHI    LCKTAKHEGALFKCDISNSTEAGQRLLQMLRLGKSEPWTLALENIVGIKTMDVKPLLNYF
W5PSB6 SHEEP        LCKTAKHEGALFKCDISNSTEAGQRLLQMLRLGKSEPWTLALENIVGIKTMDVKPLLNYF
E2DHI7 9CHIR        LCRIAQHDGPLHKCDISNSTDAGKKLHQMLSVGKSQAWTKTLEDIVDSRNMDVGPLLRYF
E2DHI4 9CHIR        LCRIAQHDGPLHKCDISNSTDAGKKLHQMLSVGKSKAWTETLKDIVDSRNMDVGPLLRYF
U5WHY8 9CHIR        LCRIAQHDGPLHKCDISNSTDAGKKLHQMLSVGKSQAWTKTLEDIVDSRNMDVGPLLKYF
G1TEF4 RABIT        LCQAAQHEGPLHKCDISNSTEAGQKLLNMLRLGRSEPWTLALENVVGAKNMDVRPLLNYF
A0A1U7QTA1 MESAU    LCQAAKHDGPLHKCDISNSTEAGQKLLNMLRLGKSEPWTLALENVVGARNMDVRPLLNYF
ACE2 MOUSE          LCQAAKYNGSLHKCDISNSTEAGQKLLKMLSLGNSEPWTKALENVVGARNMDVKPLLNYF
ACE2 RAT            LCQAAKHDGPLHKCDISNSTEAGQKLLNMLSLGNSGPWTLALENVVGSRNMDVKPLLNYF
ACE2 HUMAN          LCQAAKHEGPLHKCDISNSTEAGQKLFNMLRLGKSEPWTLALENVVGAKNMNVRPLLNYF
A0A384DV19 FELCA    LCRIAKHEGPLHKCDISNSSEAGKKLLQMLTLGKSKPWTLALEHVVGEKKMNVTPLLKYF
A0A5F5XDN9 FELCA    LCRIAKHEGPLHKCDISNSSEAGKKLLQMLTLGKSKPWTLALEHVVGEKKMNVTPLLKYF
ACE2 FELCA          LCRIAKHEGPLHKCDISNSSEAGKKLLQMLTLGKSKPWTLALEHVVGEKKMNVTPLLKYF
A0A5F4CXG9 CANLF    LCQIAKHEGPLHKCDISNSSEAGQKLLEMLKLGKSKPWTYALEIVVGAKNMDVRPLLNYF
A0A5F4BS93 CANLF    LCQIAKHEGPLHKCDISNSSEAGQKLLEMLKLGKSKPWTYALEIVVGAKNMDVRPLLNYF
F1P7C5 CANLF        LCQIAKHEGPLHKCDISNSSEAGQKLLEMLKLGKSKPWTYALEIVVGAKNMDVRPLLNYF
J9P7Y2 CANLF        LCQIAKHEGPLHKCDISNSSEAGQKLLEMLKLGKSKPWTYALEIVVGAKNMDVRPLLNYF
                    **: *:::* *.*******::**::* :** :*.*  ** :*: :*. :.*:* *** **

A0A4X1UWQ8 PIG      EPLLTWLKAQNGNSSVGWNTDWTPYADQSIKVRISLKSALGEDAYEWNDNEMYLFRSSIA
K7GLM4 PIG          EPLLTWLKAQNGNSSVGWNTDWTPYADQSIKVRISLKSALGKEAYEWNDNEMYLFRSSIA
A0A452EVU0 CAPHI    EPLFTWLKEQNRNSFVGWSTEWTPCDSVF----------VVNLQYEWNDNEMYLFRSSVA
A0A4W2H6E0 BOBOX    EPLFTWLKEQNRNSFVGWSTEWTPYSDQSIKVRISLKSALGENAYEWNDNEMYLFQSSVA
A0A4W2H3A1 BOBOX    EPLFTWLKEQNRNSFVGWSTEWTPYSDQSIKVRISLKSALGENAYEWNDNEMYLFQSSVA
Q2HJI5 BOVIN         EPLFTWLKEQNRNSFVGWSTEWTPYSDQSIKVRISLKSALGENAYEWNDNEMYLFQSSVA
ACE2 BOVIN          EPLFTWLKEQNRNSFVGWSTEWTPYSDQSIKVRISLKSALGENAYEWNDNEMYLFQSSVA
A0A452EVJ5 CAPHI    EPLFTWLKEQNRNSFVGWSTEWTPYSDQSIKVRISLKSALGENAYEWNDNEMYLFRSSVA
W5PSB6 SHEEP        EPLFTWLKEQNRNSFVGWSTEWTPYSDQSIKVRISLKSALGENAYEWNDNEMYLFRSSVA
E2DHI7 9CHIR        EPLYTWLQEQNRKSYVGWNTDWSPYSDQSIKVRISLKSALGENAYEWNDNEMYLFRSSVA
E2DHI4 9CHIR        EPLYTWLQEQNRKSYVGWNTDWSPYSDQSIKVRISLKSALGENAYEWNDNEMYLFRSSVA
U5WHY8 9CHIR        EPLYTWLQEQNRKSYVGWNTDWSPYSDQSIKVRISLKSALGENAYEWNDNEMYLFRSSVA
G1TEF4 RABIT        EPLFTWLKEQNRNSFVGWSTEWTPYADQSIKVRISLKTALGDQAYEWNDSEMYLFRSSVA
A0A1U7QTA1 MESAU    EPLSVWLKEQNKNSFVGWNTDWSPYADQSIKVRISLKSALGENAYEWDDNEMYLFRASVA
ACE2 MOUSE          QPLFDWLKEQNRNSFVGWNTEWSPYADQSIKVRISLKSALGANAYEWTNNEMFLFRSSVA
ACE2 RAT            QPLFVWLKEQNRNSTVGWSTDWSPYADQSIKVRISLKSALGKNAYEWTDNEMYLFRSSVA
ACE2 HUMAN          EPLFTWLKDQNKNSFVGWSTDWSPYADQSIKVRISLKSALGDKAYEWNDNEMYLFRSSVA
A0A384DV19 FELCA    EPLFTWLKEQNRNSFVGWNTDWRPYADQSIKVRISLKSALGDEAYEWNDNEMYLFRSSVA
A0A5F5XDN9 FELCA    EPLFTWLKEQNRNSFVGWNTDWRPYADQSIKVRISLKSALGDEAYEWNDNEMYLFRSSVA
ACE2 FELCA          EPLFTWLKEQNRNSFVGWNTDWRPYADQSIKVRISLKSALGDEAYEWNDNEMYLFRSSVA
A0A5F4CXG9 CANLF    EPLFTWLKEQNRNSFVGWNTDWSPYADQSIKVRISLKSALGEKAYEWNNNEMYLFRSSIA
A0A5F4BS93 CANLF    EPLFTWLKEQNRNSFVGWNTDWSPYADQSIKVRISLKSALGEKAYEWNNNEMYLFRSSIA
F1P7C5 CANLF        EPLFTWLKEQNRNSFVGWNTDWSPYADQSIKVRISLKSALGEKAYEWNNNEMYLFRSSIA
J9P7Y2 CANLF        EPLFTWLKEQNRNSFVGWNTDWSPYADQSIKVRISLKSALGEKAYEWNNNEMYLFRSSIA
                    :**  **: ** :* ***.*:* *  .            :    *** :.**:**::*:*

A0A4X1UWQ8 PIG      YAMRKYFSKVKNETIPFGAEDVWVSDLKPRISFNFFVTSPANMSDIIPRSDVEEAISMSR
K7GLM4 PIG          YAMRNYFSSAKNETIPFGAEDVWVSDLKPRISFNFFVTSPANMSDIIPRSDVEKAISMSR
A0A452EVU0 CAPHI    YAMRKYFLEDRNETIPFGEENVWVSDKKPRISFKFFVTSPNNVSDIIPRTEVENAIRLCR
A0A4W2H6E0 BOBOX    YAMRKYFSAARNETILFGEDNVWVSDKKPRISFKFFVTSPNNVSDIIPRTEVENAIRLSR
A0A4W2H3A1 BOBOX    YAMRKYFSAARNETILFGEDNVWVSDKKPRISFKFFVTSPNNVSDIIPRTEVENAIRLSR
Q2HJI5 BOVIN         YAMRKYFSEARNETVLFGEDNVWVSDKKPRISFKFFVTSPNNVSDIIPRTEVENAIRLSR
ACE2 BOVIN          YAMRKYFSEARNETVLFGEDNVWVSDKKPRISFKFFVTSPNNVSDIIPRTEVENAIRLSR
A0A452EVJ5 CAPHI    YAMRKYFLEDRNETIPFGEENVWVSDKKPRISFKFFVTSPNNVSDIIPRTEVENAIRLCR
W5PSB6 SHEEP        YAMRKYFLKERNETIPFGEENVWVSDKKPRISFKFFVTSPNNVSDIIPRTEVENAIRLCR
E2DHI7 9CHIR        YAMREYFLKEKHQTILFGAENVWVSNLKPRISFNFHVTSPGNLSDIIPRPEVEGAIRMSR
E2DHI4 9CHIR        YAMREYFLKEKHQTILFGAENVWVSNLKPRISFNFHVTSPGNLSDIIPRPEVEGAIRMSR
U5WHY8 9CHIR        YAMREYFLKEKHQTILFGAENVWVSNLKPRISFNFHVTSPGNLSDIIPRPEVEGAIRMSR
G1TEF4 RABIT        YAMRKYFSEVKNQTILFGEEDVRVSDLKPRISFNFFVTAPNNVNDIIPRNEVEEAISMSR
A0A1U7QTA1 MESAU    YAMRVYFAKNKTQTVPFGVEDIRVSDLKPRVSFNFFVTSPQNVSDIIPRNEVEEAVRLSR
ACE2 MOUSE          YAMRKYFSIIKNQTVPFLEEDVRVSDLKPRVSFYFFVTSPQNVSDVIPRSEVEDAIRMSR
ACE2 RAT            YAMREYFSREKNQTVPFGEADVWVSDLKPRVSFNFFVTSPKNVSDIIPRSEVEEAIRMSR
```

```
ACE2 HUMAN         YAMRQYFLKVKNQMILFGEEDVRVANLKPRISFNFFVTAPKNVSDIIPRTEVEKAIRMSR
A0A384DV19 FELCA   YAMREYFSKVKNQTIPFVEDNVWVSNLKPRISFNFFVTASKNVSDVIPRSEVEEAIRMSR
A0A5F5XDN9 FELCA   YAMREYFSKVKNQTIPFVEDNVWVSNLKPRISFNFFVTASKNVSDVIPRSEVEEAIRMSR
ACE2 FELCA         YAMREYFSKVKNQTIPFVEDNVWVSNLKPRISFNFFVTASKNVSDVIPRSEVEEAIRMSR
A0A5F4CXG9 CANLF   YAMRQYFSEVKNQTIPFVEDNVWVSDLKPRISFNFFVTSPGNVSDIIPRTEVEEAIRMYR
A0A5F4BS93 CANLF   YAMRQYFSEVKNQTIPFVEDNVWVSDLKPRISFNFFVTSPGNVSDIIPRTEVEEAIR---
F1P7C5 CANLF       YAMRQYFSEVKNQTIPFV---------------------------------------MYR
J9P7Y2 CANLF       YAMRQYFSEVKNQTIPFVEDNVWVSDLKPRISFNFFVTSPGNVSDIIPRTEVEEAIRMYR
                   **** **   : : : *

A0A4X1UWQ8 PIG     SRINDAFRLDDNTLEFLGIQPTLGPPDEPPVTVWLIIFGVVMGLVVVGIVVLIFTGIRDR
K7GLM4 PIG         SRINDAFRLDDNTLEFLGIQPTLGPPDEPPVTVWLIIFGVVMGLVVVGIVVLIFTGIRDR
A0A452EVU0 CAPHI   DRINDAFQLDDNSLEFLGIQPTLRPPYEPPVTIWLIIFGVVMGVVVIGIVVLIFTGIRDQ
A0A4W2H6E0 BOBOX   DRINDVFQLDDNSLEFLGIQPTLGPPYEPPVTIWLIIFGVVMGVVVIGIVVLIFTGIRNR
A0A4W2H3A1 BOBOX   DRINDVFQLDDNSLEFLGIQPTLGPPYEPPVTIWLIIFGVVMGVVVIGIVVLIFTGIRNR
Q2HJI5 BOVIN       DRFNDVFQLDDNSLEFLGIQPTLGPPYEPPVTIWLIIFGVVMGVVVIGIVVLIFTGIRNR
ACE2 BOVIN         DRINDVFQLDDNSLEFLGIQPTLGPPYEPPVTIWLIIFGVVMGVVVIGIVVLIFTGIRNR
A0A452EVJ5 CAPHI   DRINDAFQLDDNSLEFLGIQPTLRPPYEPPVTIWLIIFGVVMGVVVIGIVVLIFTGIRDQ
W5PSB6 SHEEP       DRINDAFQLDDNSLEFLGIQPTLRPPYEPPVTIWLIIFGVVMGVVVIGIVVLIFTGIRDQ
E2DHI7 9CHIR       SRINDAFRLDDNSLEFLGIQPTLGPPYQPPVTIWLIVFGVVMAVVVVGIVVLIITGIRDR
E2DHI4 9CHIR       SRINDAFRLDDNSLEFLGIQPTLGPPYQPPVTIWLIVFGVVMAVVVVGIVVLIITGIRDR
U5WHY8 9CHIR       SRINDAFRLDDNSLEFLGIQPTLGPPYQPPVTIWLIVFGVVMAVVVVGIVVLIITGIRDR
G1TEF4 RABIT       SRINDIFRLDDNSLEFVGIQPTLEPPYESPVPIWLVVFGVVMGMIVIGIVVLIFTGIKDR
A0A1U7QTA1 MESAU   GRINDVFGLDDNSLEFLGINPTLSPPYQPPVTIWLIIFGVVMGIVVVGIIILIFTGIKGR
ACE2 MOUSE         GRINDVFGLNDNSLEFLGIHPTLEPPYQPPVTIWLIIFGVVMALVVVGIIILIVTGIKGR
ACE2 RAT           GRINDIFGLNDNSLEFLGIYPTLKPPYEPPVTIWLIIFGVVMGTVVVGIVILIVTGIKGR
ACE2 HUMAN         SRINDAFRLNDNSLEFLGIQPTLGPPNQPPVSIWLIVFGVVMGVIVVGIVILIFTGIRDR
A0A384DV19 FELCA   SRINDAFRLDDNSLEFLGIQPTLSPPYQPPVTIWLIVFGVVMGVVVVGIVLLIVSGIRNR
A0A5F5XDN9 FELCA   SRINDAFRLDDNSLEFLGIQPTLSPPYQPPVTIWLIVFGVVMGVVVVGIVLLIVSGIRNR
ACE2 FELCA         SRINDAFRLDDNSLEFLGIQPTLSPPYQPPVTIWLIVFGVVMGVVVVGIVLLIVSGIRNR
A0A5F4CXG9 CANLF   SRINDVFRLDDNSLEFLGIQPTLGPPYEPPVTIWLIVFGVVMGVVVVGIVLLIFSGIRNR
A0A5F4BS93 CANLF   ------------------------------------------------------------
F1P7C5 CANLF       SRINDVFRLDDNSLEFLGIQPTLGPPYEPPVTIWLIVFGVVMGVVVVGIVLLIFSGIRNR
J9P7Y2 CANLF       SRINDVFRLDDNSLEFLGIQPTLGPPYEPPVTIWLIVFGVVMGVVVVGIVLLIFSGIRNR


A0A4X1UWQ8 PIG     RKKKQASSEENPYGSMDLSK---------GESNSGFQNGDDIQTSF
K7GLM4 PIG         RKKKQASSEENPYGSMDLSK---------GESNSGFQNGDDIQTSF
A0A452EVU0 CAPHI   RKKNQASSEENPYGSVD-------LNKG--ENNSGFQNTDDVQTSL
A0A4W2H6E0 BOBOX   RKHDSDGLQNDENLRVQQQAVKVDIPRNSLKATVPFSNSHEKLK--
A0A4W2H3A1 BOBOX   RKKNQASSEENPYGSVD-------LNKGE--NNSGFQNIDDVQTSL
Q2HJI5 BOVIN       RKKNQASSEENPYGSVD-------LNKGE--NNSGFQNIDDVQTSL
ACE2 BOVIN         RKKNQASSEENPYGSVD-------LNKGE--NNSGFQNIDDVQTSL
A0A452EVJ5 CAPHI   RKKNQASSEENPYGSVD-------LNKGE--NNSGFQNTDDVQTSL
W5PSB6 SHEEP       RKKNQASSEENPYGSVD-------LNKGE--NNSGFQNTDDVQTSL
E2DHI7 9CHIR       RKTDQARSEENPYSSVDLSK---------GENNPGFQNGDDVQTSF
E2DHI4 9CHIR       RKTDQARSEENPYSSVDLSK---------GENNPGFQNGDDVQTSF
U5WHY8 9CHIR       RKTDQARSEENPYSSVDLSK---------GENNPGFQNGDDVQTSF
G1TEF4 RABIT       RKQKQAKREENPYGFVDMSK---------GENNSGFQNSDDIQTSF
A0A1U7QTA1 MESAU   KKKNETKREENPYDSVDIGK---------GESNAGFLSNDDAQTSF
ACE2 MOUSE         KKKNETKREENPYDSMDIGK---------GESNAGFQNSDDAQTSF
ACE2 RAT           KKKNETKREENPYDSMDIGK---------GESNAGFQNSDDAQTSF
ACE2 HUMAN         KKKNKARSGENPYASIDISK---------GENNPGFQNTDDVQTSF
A0A384DV19 FELCA   RKNNQARSEENPYASVDLSK---------GENNPGFQHADDVQTSF
A0A5F5XDN9 FELCA   RKNNQARSEENPYASVDLSK---------GENNPGFQHADDVQTSF
ACE2 FELCA         RKNNQARSEENPYASVDLSK---------GENNPGFQHADDVQTSF
A0A5F4CXG9 CANLF   RKWPSSLSSP------------------------------------
A0A5F4BS93 CANLF   --NDQARGEENPYASVDLSK---------GENNPGFQNVDDAQTSF
F1P7C5 CANLF       RKNDQARGEENPYASVDLSK---------GENNPGFQNVDDAQTSF
J9P7Y2 CANLF       RKNDQARGEENPYASVDLSK---------GENNPGFQNVDDAQTSF
```
